# Tyr1 phosphorylation promotes the phosphorylation of Ser2 on the C-terminal domain of RNA polymerase II by P-TEFb

**DOI:** 10.1101/652214

**Authors:** Joshua E. Mayfield, Seema Irani, Edwin E. Escobar, Zhao Zhang, Nathanial T. Burkholder, Michelle R. Robinson, M. Rachel Mehaffey, Sarah N. Sipe, Wanjie Yang, Nicholas A. Prescott, Karan R. Kathuria, Zhijie Liu, Jennifer S. Brodbelt, Yan Zhang

## Abstract

The Positive Transcription Elongation Factor b (P-TEFb) phosphorylates Ser2 residues of RNA polymerase II’s C-terminal domain (CTD) and is essential for the transition from transcription initiation to elongation *in vivo*. Surprisingly, P-TEFb exhibits Ser5 phosphorylation activity *in vitro*. The mechanism garnering Ser2 specificity to P-TEFb remains elusive and hinders understanding of the transition from transcription initiation to elongation. Through *in vitro* reconstruction of CTD phosphorylation, mass spectrometry analysis, and chromatin immunoprecipitation sequencing (ChIP-seq) analysis we uncover a mechanism by which Tyr1 phosphorylation directs the kinase activity of P-TEFb and alters its specificity from Ser5 to Ser2. The loss of Tyr1 phosphorylation causes a reduction of phosphorylated Ser2 and accumulation of RNA polymerase II in the promoter region as detected by ChIP-seq. We demonstrate the ability of Tyr1 phosphorylation to generate a heterogeneous CTD modification landscape that expands the CTD’s coding potential. These findings provide direct experimental evidence for a combinatorial CTD phosphorylation code wherein previously installed modifications direct the identity and abundance of subsequent coding events by influencing the behavior of downstream enzymes.

---

The C-terminal domain of RNA polymerase II (CTD) is composed of a species-specific number of repeats of the consensus amino acid heptad YSPTSPS (arbitrarily numbered as Tyr1, Ser2, Pro3, Thr4, Ser5, Pro6, & Ser7) (Jeronimo et al., 2016). The CTD undergoes extensive post-translational modification (PTM) that recruits RNA processing and transcription factors that regulate progression through the various stages of transcription. These modification events are dynamic, highly regulated, and maintained through the complex interplay of CTD modification enzymes. Collectively these PTMs and recruited protein factors constitute the “CTD Code” for eukaryotic transcription (Buratowski, 2003).

Chromatin immunoprecipitation and next-generation sequencing technologies have revealed how phosphorylation levels of CTD residues change temporally and spatially during each transcription cycle (Eick and Geyer, 2013). The major sites of phosphorylation are Ser5, directed by TFIIH in mammals (Feaver et al., 1994), and Ser2, installed by P-TEFb in mammals (Marshall et al., 1996). The other three phosphate-accepting residues (Tyr1, Thr4, and Ser7) are also subject to modification, although their functions are less well-understood (Jeronimo et al., 2013). In mammalian cells, the phosphorylations of Tyr1 and Ser7 rise and peak near the promoter along with Ser5 and gradually decrease as transcription progresses towards termination. The phosphorylation of Thr4 and Ser2, on the other hand, don’t appear until later in the transcription cycle during elongation (Eick and Geyer, 2013). The molecular underpinnings resulting in this orchestration are poorly defined. A particularly apparent gap in current knowledge is if sequence divergence from the consensus heptad or previously installed PTMs influence coding events.

The CTD code is generated through the interplay of CTD modifying enzymes such as kinases, phosphatases and prolyl isomerases (Bataille et al., 2012). Disruption of this process is implicated in various disease states. P-TEFb is of particular interest due to its overexpression in multiple tumor types and role in HIV infection (Franco et al., 2018). As a major CTD kinase, P-TEFb promotes transcription by contributing to the release of RNA polymerase II from the promoter-proximal pause through its phosphorylation of Negative Elongation Factor (NELF), DRB Sensitivity Inducing Factor (DSIF), and Ser2 of the CTD (Wada et al., 1998). Interestingly, P-TEFb seems to phosphorylate Ser5 of the CTD *in vitro* and mutation of Ser5 to alanine prevents the phosphorylation of CTD substrates. However, mutation of Ser2 to alanine did not result in this abolishment (Czudnochowski et al., 2012). These results are in contrast to *in vivo* studies of P-TEFb specificity, where compromised P-TEFb kinase activity results in a specific reduction in levels of Ser2 phosphorylation (Marshall et al., 1996). The discrepancies between P-TEFb specificity *in vitro* and *in vivo* make it difficult to reconcile P-TEFb’s function as a CTD Ser2 kinases (Bartkowiak et al., 2010; Czudnochowski et al., 2012).

To resolve these inconsistencies, we utilize a multi-disciplinary approach to investigate the specificity of P-TEFb. Identification of phosphorylation sites was carried out using ultraviolet photodissociation (UVPD) mass spectrometry establishing the specificity of P-TEFb *in vitro* with single residue resolution. We reveal the tyrosine kinase c-Abl phosphorylates consensus and full-length CTD substrates in a conservative fashion, with only half of the available sites phosphorylated. The unique phosphorylation pattern of Tyr1 by tyrosine kinases like c-Abl directs the specificity of P-TEFb to Ser2. The priming effect of pTyr1 on P-TEFb extends to human cells, where small molecule inhibition of c-Abl-like Tyr1 kinase activity leads to a reduction of Tyr1 phosphorylation accompanied by a significant decrease in Ser2 phosphorylation. Further ChIP-seq analysis shows that the loss of tyrosine phosphorylation increases promoter-proximal pausing with an accumulation of RNA polymerase II and a reduction of pSer2 in the promoter region of the genes. Overall, our results reconcile the discrepancy of P-TEFb kinase activity *in vitro* and in cells, showing that Tyr1 phosphorylation can prime P-TEFb and alter its specificity to Ser2 to overcome promoter-proximal pausing during eukaryotic transcription. Importantly, these findings provide direct experimental evidence for a combinatorial CTD phosphorylation code wherein previously installed modifications direct the identity and abundance of subsequent coding events, resulting in a varied PTM landscape along the CTD allowing for diversified co-transcriptional signaling.

## Results

### Determination of the in vitro P-TEFb specificity using mass spectrometry

To define P-TEFb specificity directly on endogenous CTD of RNA polymerase II we applied matrix assisted laser desorption/ionization mass spectrometry (MALDI-MS) and liquid chromatography ultraviolet photodissociation tandem mass spectrometry (LC-UVPD-MS/MS) to identify the substrate residues of this kinase. Because endogenous RNA polymerase II is highly heterogeneously modified, we used recombinant yeast CTD GST fusion proteins, which contain mostly consensus heptad repeats (20 of 26), as an unmodified substrate for PTM analysis (Figure S1A). The stability and consistency of GST yeast CTD (yCTD) make it ideal for studying CTD modification along consensus heptads. Under saturating conditions, P-TEFb generates two phospho-peptides as detected by LC-UVPD-MS/MS: a major species phosphorylated on Ser5 (Y_1_S_2_P_3_T_4_**<u>pS</u>**_5_P_6_S_7_) and a minor species phosphorylated on Ser2 (S_5_P_6_S_7_Y_1_**<u>pS</u>**_2_P_3_T_4_) (Figure 1A and S2A-B). This is highly similar to patterns observed previously for *bona fide* Ser5 CTD kinases Erk2 and TFIIH (Mayfield et al., 2017). These experiments confirm P-TEFb’s inherent *in vitro* preference for Ser5 when phosphorylating unmodified CTD (Czudnochowski et al., 2012; Portz et al., 2017).

**Figure 1:**
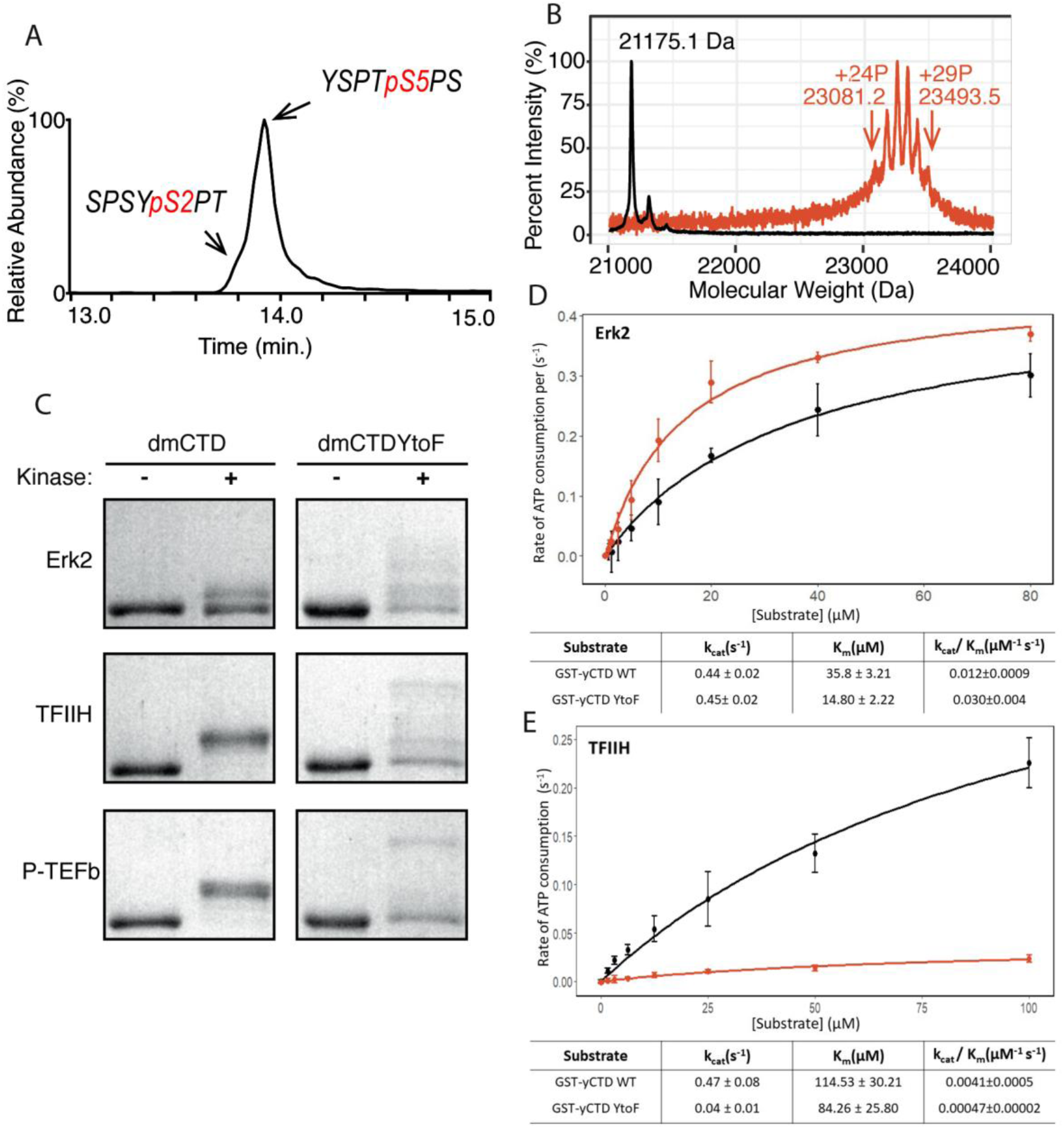
P-TEFb in vitro activity and phenylalanine replacement of Tyr1 alters phosphorylation of the CTD. (**A**) LC-UVPD-MS/MS analysis of yCTD treated with P-TEFb alone showing extracted ion chromatogram for two CTD heptads. (**B**) Portions of MALDI mass spectra of 3C-protease digested yCTD construct treated with P-TEFb alone (red) and no kinase control reaction (black). Mass labels indicate m/z at the various peak maxima. Arrows indicate the maximum and minimum m/z peaks for kinase treated sample. “+#P” notation indicates an approximate number of phosphates added based on mass shifts relative to no kinase control. (**C**) SDS-PAGE EMSA of dmCTD and dmCTDYtoF (as indicated) treated with Erk2 (top, right bands), TFIIH (middle, right bands), or P-TEFb (bottom, right bands) and paired no kinase control reactions (left bands). Biological triplicate were carried out with three different reactions of same reaction were setup. (**D-E**) Kinase activity assay of wild-type yCTD (shown in black) and yCTDYtoF (shown in red) variant by Erk2 (**D**) and TFIIH (**E**) fitted to the Michaelis-Menten kinetic equation. The Michaelis-Menten kinetic parameters *k_cat_*(s^-1^), *K_m_*(µM) and *k_cat_/K_m_*(µM^-1^ s^-1^) are given below the graphs for each respective fit. Each measurement was conducted in triplicate (technical triplicate) with standard deviations shown as error bars.

We next measured the total number of phosphates added to CTD by P-TEFb. MALDI-MS analysis of yCTD treated with P-TEFb reveals a cluster of peaks with mass shifts relative to no kinase control ranging from 1906.1 to 2318.4 Da, each interspaced by 80 Da (Figure 1B, S2C). This corresponds to the addition of 24 to 29 phosphates along yCTD’s 26 heptad repeats. This finding in combination with our LC-UVPD-MS/MS analysis of P-TEFb treated yCTD indicates that P-TEFb phosphorylates the CTD in a one phosphorylation per heptad manner and these heptads are primarily phosphoryl-Ser5 (pSer5) *in vitro*.

### Amino acid identity at the Tyr1 position is important for kinase activity

To phosphorylate Ser2 and Ser5, CTD kinases must discriminate very similar SP motifs in the CTD, Y_1_**<u>S_2_</u>**P_3_ and T_4_**<u>S_5_</u>**P_6_, to maintain accuracy during transcription. Among the flanking residues of these two motifs, the unique structure of the tyrosine side chain likely contributes to the recognition of the serine residues subject to phosphorylation. Several factors suggest the chemical properties of residues at the Tyr1 position are important for CTD modification. First, residues at this first position of the heptad are highly conserved across species and substitution to non-aromatic residues is rare, suggesting significance to function (Chapman et al., 2008). As evidence of this, even conservative mutation of the Tyr1 position to phenylalanine in both *Saccharomyces cerevisiae* and human cells is lethal and highlights the significance of residues at this position (Hsin et al., 2014; West and Corden, 1995). Secondly, we have shown that mutating the Tyr1 position to alanine prevents phosphorylation at other CTD residues by CTD kinases (Mayfield et al., 2017), indicating the side chain at this position is important for kinase activity. Third, phosphoryl-Tyr1 (pTyr1) is detected at the initiation of transcription in human cells (Descostes et al., 2014). This positions pTyr1 well to influence and interact with subsequent modifications of the CTD and potentially influence subsequent enzyme specificity.

To determine the effect of the chemical characteristics of residues at the Tyr1 position on CTD modification, we searched for naturally occurring Tyr1 substitutions. *Drosophila melanogaster* CTD contains a majority of heptads that diverge from consensus sequence with only 2 of its approximately 45 heptads being of the consensus sequence. Despite this highly divergent character, the Tyr1 position of *D. melanogaster* CTD is strongly conserved and contains mostly tyrosine residues. For the six heptads that do not contain tyrosine, half are modestly substituted with a phenylalanine. We were curious to determine if, like alanine, phenylalanine replacement at the Tyr1 position would abolish CTD kinase activity. We generated GST-CTD constructs containing a nearly consensus portion of *D. melanogaster* CTD (residues 1671-1733, containing nine heptad repeats) of either tyrosine containing wild-type (dmCTD) or with phenylalanine substitution at the Tyr1 position in all nine heptads (dmCTDYtoF) (Figure S1D). These constructs were phosphorylated with one of three established CTD kinases: Erk2, a recently identified Ser5 CTD kinase that phosphorylates primed RNA polymerase II in developmental contexts (Tee et al., 2014); the kinase module of TFIIH that install Ser5 and Ser7 marks *in vivo* (Feaver et al., 1994); or P-TEFb which phosphorylates Ser2 *in vivo* (Marshall et al., 1996). Unlike alanine substitution, all three kinases are active against the phenylalanine-substituted CTD construct. Surprisingly, substitution of phenylalanine at the Tyr1 position alters the behavior of phosphorylated substrates in electrophoretic mobility shift assays (EMSA) (Figure 1C). While the wild-type variant assumes only one or two apparent intermediate species in EMSA, the YtoF variant of dmCTD exhibits multiple intermediates suggesting the generation of a greater diversity of phosphorylated species. Additional analysis of Erk2 phosphorylated dmCTDYtoF using electrospray ionization mass spectrometry (ESI-MS) of the intact phosphorylated construct confirms the existence of multiple species revealing complex spectra composed of multiple overlapping peaks relative to the dmCTD control (Figure S2D). To quantify the effect of phenylalanine replacement at the Tyr1 position on CTD kinase function we measured the kinase activity of Erk2 and TFIIH using GST-yCTD or yCTDYtoF substrate (Figure S1A-B). Steady-state kinetics demonstrate that the replacement of tyrosine by phenylalanine has a markedly different effect on these two kinases. Erk2 shows a 2.5-fold higher specificity constant against the YtoF variant, as indicated by *k_cat_*/*K_m_*, compared to the WT construct (Figure 1D). Erk2 has nearly identical *k_cat_*values (0.44 ± 0.02 s^-1^ vs. 0.45 ± 0.02 s^-1^) for the two substrates, but a much lower *K_m_*for the YtoF substrate (35.8 ± 3.2 µM for WT vs. 14.8 ± 2.2 µM for YtoF substrates). This difference in *K_m_*values suggests Erk2 has a binding preference for the phenylalanine substituted substrate. However, TFIIH activity is greatly compromised when Tyr1 is replaced by phenylalanine with a nearly 10-fold reduction in *k_cat_*/*K_m_*(Figure 1E).

Overall, our data demonstrate the chemical properties of the residue located at the first position of the heptad repeat has a significant impact on the phosphorylation of the CTD residues by CTD kinases. Even slight modification of this residue (e.g., loss of the hydroxyl group) can have dramatic consequences for modification of the CTD.

### Tyr1 phosphorylation in human CTD

Although substitution of non-tyrosine residues at the Tyr1 position is relatively rare in nature and does not occur in human cells, Tyr1 phosphorylation is conserved from yeast to humans and plays key role in transcriptional events although the molecular mechanism explaining the diverse biological functions remains elusive (Chapman et al., 2008). The sensitivity of CTD kinases to the chemical properties of Tyr1 side chain motivated us to investigate if Tyr1 phosphorylation impacts subsequent phosphorylation events by reconstructing sequential CTD phosphorylation *in vitro*. In humans, Tyr1 phosphorylation rises along with Ser5 phosphorylation at the beginning of transcription (Heidemann et al., 2013). However, experiments using synthetic CTD peptides with every Tyr1 residue phosphorylated have shown that Tyr1 phosphorylation inhibits subsequent phosphorylation by CTD kinases (Czudnochowski et al., 2012). We suspect that the heavily phosphorylated synthetic peptide doesn’t mimick the physiological RNA polymerase II during transcription well. Instead, we recapitulated the phosphorylation of the CTD using physiologically relevant Tyr1 kinases *in vitro*. Existing literature points to Abl-like non-receptor tyrosine kinases as mammalian Tyr1 CTD kinases, with c-Abl as a major candidate (Baskaran et al., 1997; Burger et al., 2019). Three lines of evidence support this notion: c-Abl phosphorylates CTD *in vitro* (Baskaran et al., 1993) and in cells since transient over-expression of c-Abl in primate COS cells results in increased Tyr1 phosphorylation (Baskaran et al., 1997), and c-Abl immunoprecipitates with RNA polymerase II (Baskaran et al., 1999). To elucidate the biophysical consequences of Tyr1 phosphorylation of the CTD we reconstructed c-Abl phosphorylation of consensus sequence CTD *in vitro* using purified human c-Abl and the yCTD constructs. C-Abl readily phosphorylates yCTD *in vitro* as evidenced by EMSA and detection of Tyr1 phosphorylation using pTyr1 specific antibody 3D12 (Figure 2A). We directly interrogate the sites of phosphorylation using LC-UVPD-MS/MS to identify phosphorylation sites in single residue resolution. Previously, we developed UVPD-MS/MS analysis strategies to investigate challenging targets such as CTD, which provides improved characterization and more diagnostic spectra (Mayfield et al., 2017). Using this method to analyze a peptide containing three heptad repeats treated by c-Abl (3CTD, Figure S1E), two single phospho-forms were detected, each containing a single phosphorylated tyrosine on either the first or second heptad of the variant (Figure 2B and S2E). These mass shifts confirm c-Abl phosphorylates consensus CTD sequences on Tyr1 *in vitro*.

**Figure 2:**
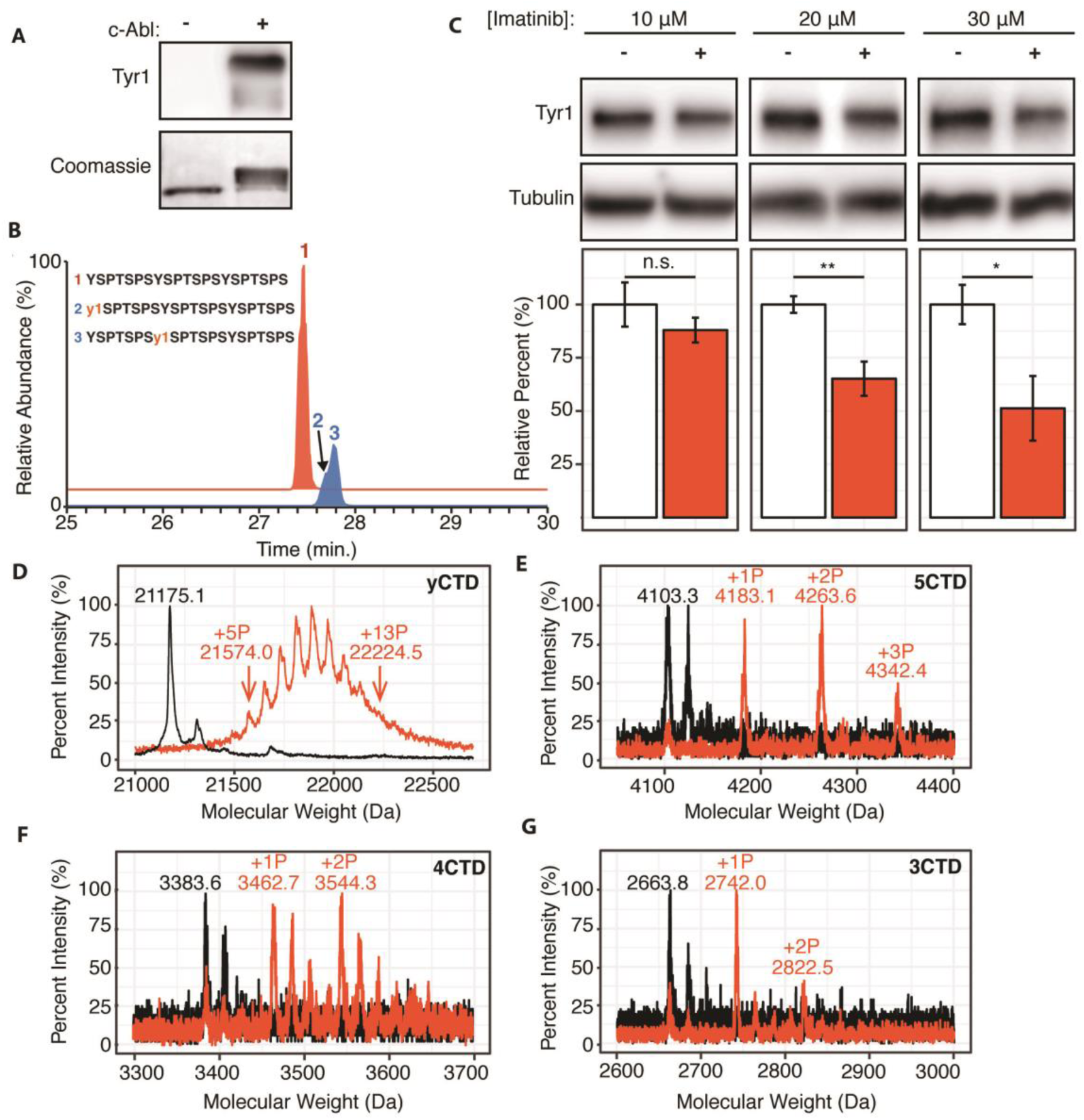
c-Abl kinase phosphorylates Tyr1 of RNA polymerase II CTD in cells and *in vitro*. (**A**) Representative image of western blot against phosphorylated Tyr1 (top) of yCTD (containing 26 heptad repeats) treated *in vitro* with c-Abl (right) and paired no kinase control (left). Coomassie-stained blot included indicating loading (bottom). Data representative of three experimental replicates. (B) LC-UVPD-MS/MS analysis of 3CTD treated with c-Abl showing extracted ion chromatograms for 3CTD (m/z 888.76, 3+ charge state, red XIC) and mono-phosphorylated 3CTD (m/z 915.36, 3+ charge state, blue XIC) peptides. **(C)** Representative images (top) and quantification (bottom) of a western blot of imatinib dosage series (10-30 μM, as indicated, red) and paired DMSO vehicle controls (left band, white) of 20 μg total protein from HEK293T cells. Phospho-specific Tyr1 antibody (clone 3D12) was used. Imatinib decreases pTyr1 epitope abundance to 88.0% (10μM imatinib, not significant), 65.2% (20μM imatinib), and 51.3% (30μM imatinib) relative to paired vehicle controls. Epitope signals normalized against tubulin loading control. Significance determined by Welch’s t-test (* = p-value < 0.05, ** = p-value < 0.01, n.s. = not significant (p-value > 0.05)), n=6, error bars indicate SEM. (**D-G**) Portions of MALDI mass spectra of 3C-protease digested yeast CTD (**D**), 5CTD (**E**), 4CTD (**F**), and 3CTD (**G**) construct treated with c-Abl (red) and no kinase control reaction (black). Mass labels indicate m/z at various peak maxima. Arrows indicate the maximum and minimum m/z peak for kinase treated sample. “+#P” notation indicates an approximate number of phosphates added based on mass shifts. Satellite peaks, prevalent in (E), correlate well with sodium adducts (M+23 Da).

To test if modulating c-Abl activity can alter Tyr1 phosphorylation levels in human cells, we treated HEK293T cells with the c-Abl specific inhibitor imatinib (Knight and McLellan, 2004) and monitored endogenous Tyr1 phosphorylation levels using phospho-Tyr1 specific antibody 3D12 (Figure 2C). Tyr1 phosphorylation decreases in a dose-dependent manner from 10-50% at imatinib concentrations of 10-30 μM after 24 hours of treatment (Figure 2C). Overall, our result indicates controlling the kinase activity of c-Abl, or highly similar kinases, is sufficient to significantly modulate the level of Tyr1 phosphorylation of CTD in mammalian cells.

We next quantified the maximal number of phosphates added to yCTD constructs using MALDI-MS. High-resolution MALDI-MS spectra of samples treated by c--Abl under saturating conditions revealed peaks accounting for yCTD containing 5 to 13 phosphates (with mass shifts ranging from 398.9 to 1049.4 Da) (Figure 2D), approximately half of yCTD’s available tyrosine residues within 26 heptad repeats. Additional GST-CTD constructs containing 3-5 consensus heptad repeats (Figure S1E-G) treated with c-Abl under saturating conditions were measured using MALDI-TOF to evaluate if c-Abl truly only phosphorylates half of the available Tyr1 sites. Three phosphorylation peaks were detected in the 5CTD variant with mass differences of 79.8, 160.3 and 239.1 Da relative to the unphosphorylated control, accounting for the addition of 1-3 phosphates (Figure 2E). Phosphorylation of the 4CTD construct resulted in two peaks of phosphorylation with mass differences of 79.1 and 160.7 Da relative to unphosphorylated control, accounting for the addition of 1 or 2 phosphates (Figure 2F). Similarly, two phosphates are added to the 3CTD variant that displayed mass shifts of 78.2 or 158.7 Da (Figure 2G). These mass shifts suggest c-Abl does not phosphorylate consensus CTD in every heptad instead it favors phosphorylation of approximately half the available Tyr1 residues.

### Sequential phosphorylation of CTD by c-Abl followed by P-TEFb

With our knowledge of the previously undescribed Tyr1 phosphorylation pattern installed by c-Abl, we were curious if such a pattern could affect the phosphorylation of CTD by P-TEFb. We first determined if the pre-treatment of CTD by c-Abl alters the number of phosphates added by P-TEFb using MALDI-TOF. Since c-Abl phosphorylates tyrosine and P-TEFb phosphorylates serine residues as determined (Figure 2A and 1A, respectively), if the two phosphorylation events are independent, the phosphate placed by the kinases should be the addition of the two kinases individually. C-Abl phosphorylation of yCTD alone adds up to 13 phosphates (Figure 2D) and P-TEFb alone adds 24-29 phosphates (Figure 1B). Interestingly, tandem treatment of yCTD with c-Abl followed by P-TEFb resulted in the addition of a total of 16 to 26 phosphates as detected by MALDI-MS, with a mass shifts of 1287.3 to 2073.2 Da (Figure 3A). This data reveals c-Abl pre-treatment results in changes to P-TEFb’s phosphorylation along the CTD, evidenced by a reduction in the number of phosphate groups added by P-TEFb.

**Figure 3:**
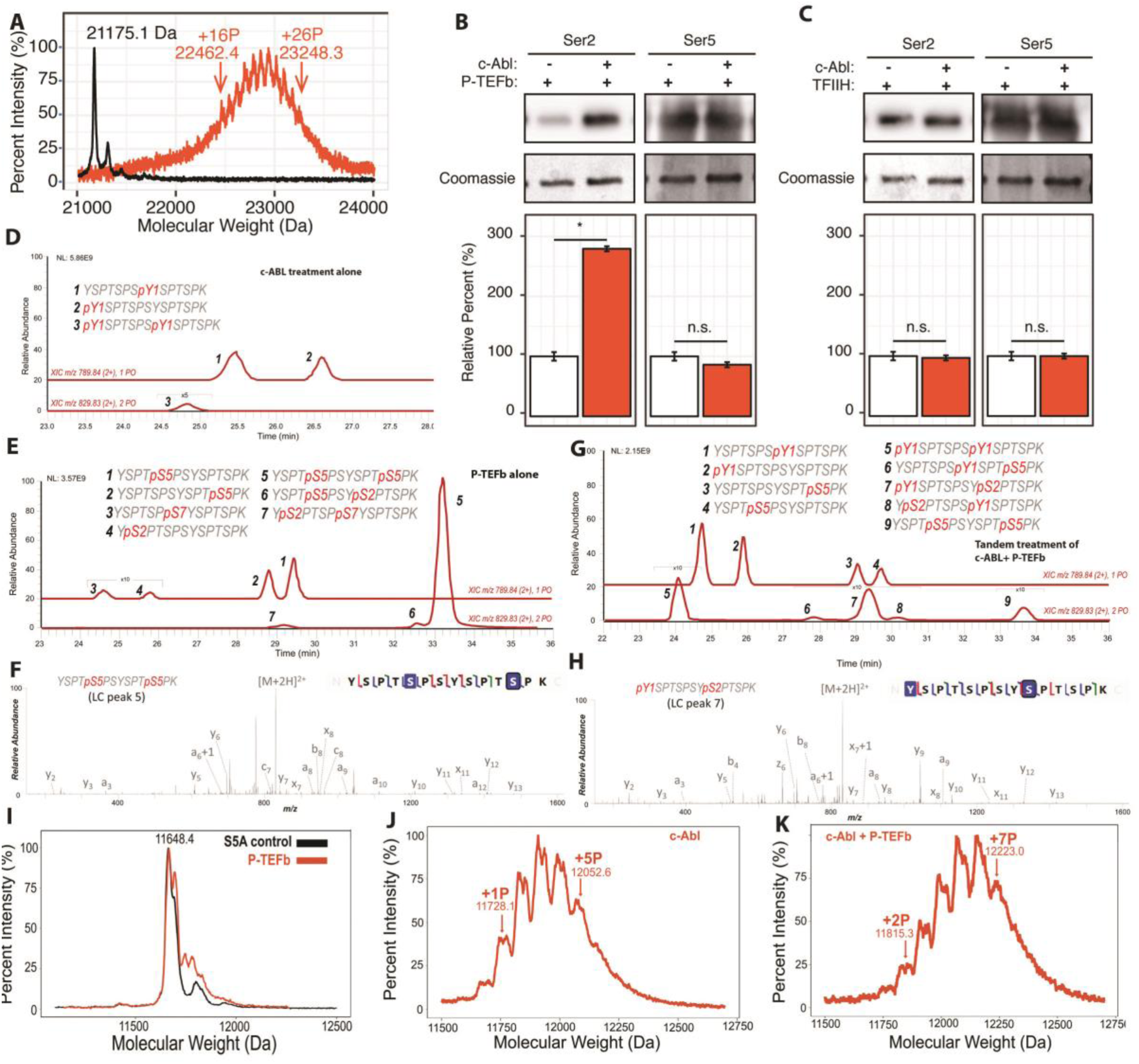
Effect of Tyr1 phosphorylation by c-Abl on the function of P-TEFb. (**A**) Portions of MALDI mass spectra of 3C-protease digested yCTD construct treated tandemly with c-Abl followed by P-TEFb (red) and no kinase control reaction (black). Mass labels indicate m/z peaks for kinase treated sample. “+#P” notation indicates an approximate number of phosphates added based on mass shifts. (**B**) Representative images (top) and quantification (bottom) of western blot analysis of yCTD treated with P-TEFb alone (left, white) and tandemly with c-Abl followed by P-TEFb (right, red). Tandem treatment of c-Abl followed by P-TEFb increases phosphorylated Ser2 epitope abundance to 279% of P-TEFb only treatment control. Ser5 phosphorylation levels is not significantly altered. Significance determined by Welch’s t-test (* = p-value < 0.05, n.s. = not significant (p-value > 0.05)), n = 3, error bars indicate SEM. (**C**) Representative images (top) and quantification (bottom) of western blot analysis of yCTD treated with TFIIH alone (left, white) and tandemly with c-Abl followed by TFIIH (right, red). Tandem treatment of c-Abl followed by TFIIH does not significantly alter the epitope abundance of phosphorylated Ser2 or Ser5. Significance determined by Welch’s t-test (n.s. = not significant (p-value > 0.05)), n = 3, error bars indicate SEM. (**D-H**) LC-UVPD-MS/MS analysis of yeast CTD with inserted Lys in every other heptad repeat (yCTD-Lys) treated with kinases or combination of kinases. Biological triplet samples were independently measured with exemplary spectra shown (n=3). (**D**) yCTD-Lys treated with c-Abl showing extracted ion chromatograms for mono-phosphorylated (m/z 789.84, 2+ charge state) and doubly phosphorylated (m/z 829.83, 2+ charge state) peptides of sequence (YSPTSPSYSPTSPK). The LC traces are shown in red and the phosphorylation sites determined by UVPD-MS/MS are highlighted in red font with “p” to indicate phosphorylation. (**E**) LC-UVPD-MS/MS analysis of yCTD-Lys treated with P-TEFb showing extracted ion chromatograms for mono-phosphorylated (m/z 789.84, 2+ charge state) and doubly phosphorylated (m/z 829.83, 2+ charge state) peptides. (**F**) Representative UVPD spectra that demonstrate the diagnostic fragmentation pattern of the peptides shown in the inset from (E). The one shown is peak 5, which is the predominant product from yCTD-Lys treated by P-TEFb alone. (**G**)LC-UVPD-MS/MS analysis of yCTD-Lys with c-Abl followed by P-TEFb showing extracted ion chromatograms for mono-phosphorylated (m/z 789.84, 2+ charge state) and doubly phosphorylated (m/z 829.83, 2+ charge state) peptides. (**H**) Representative UVPD spectra that demonstrate the diagnostic fragmentation pattern of the peptides shown in the inset of (G). The one shown is peak 7, which is the predominant product from yCTD-Lys treated by c-Abl followed by P-TEFb. (**I-K**) Portions of MALDI-MS spectra of 3C-digested S5A. Panels included no kinase control (**I**), P-TEFb only treated (**J**), c-Abl only treated (**I**), and tandemly treated with c-Abl followed by P-TEFb (**K**). Mass labels indicate m/z at the various peak maxima. Arrows indicate the maximum and minimum m/z peak for kinase treated sample. “+#P” notation indicates an approximate number of phosphates added based on mass shifts.

To identify the position of phosphates added by P-TEFb when Tyr1 is phosphorylated, we quantified pSer2 and pSer5 by immunoblotting with antibodies recognizing Ser2 and Ser5 phosphorylations (Figure 3B). Compared to a non-phosphorylated CTD, the pre-treatment of yCTD with c-Abl results in a significant increase in Ser2 phosphorylation of nearly 300% as detected by pSer2 specific CTD antibody 3E10, accompanied by little change in pSer5 by P-TEFb (Figure 3B and 3C). The increase of pSer2 is unique for P-TEFb-mediated phosphorylation on CTD since similar tandem treatment of c-Abl followed by either TFIIH or Erk2 showed no changes on pSer2 levels (Figure S3A).

We propose two possible explanations for the apparent increase of pSer2 levels upon c-Able/P-TEFb treatment: First, c-Abl interacts with and/or modifies P-TEFb and alters its specificity from Ser5 to Ser2. Alternatively, c-Abl may phosphorylate substrate CTD and these phosphorylations prime the P-TEFb specificity towards Ser2 residues of the CTD. To differentiate these two models, we inactivated c-Abl after its reaction with CTD but before the addition of P-TEFb. We used two independent methods to inactivate c-Abl prior to P-TEFb addition: introduction of the potent Abl inhibitor dasatinib to 10 µM or denaturation of c-Abl via heat-inactivation (Figure S3B). In both experiments, P-TEFb continues to install a greater amount of Ser2 phosphorylation relative to no c-Abl treatment controls (Figure S3B). Therefore, the increase in the apparent Ser2 phosphorylation is not due to P-TEFb’s physical interaction with c-Abl but arises from c-Abl kinase activity against CTD substrates at Tyr1.

To determine the phosphorylation pattern resulting from sequential kinase treatment, we used LC-UVPD-MS/MS to investigate the activity of P-TEFb in the context of Tyr1 phosphorylation. LC-UVPD-MS/MS provides single residue resolution and overcomes artifacts inherent to immunoblotting such as epitope masking. Unfortunately, full-length yeast CTD is resistant to proteolysis due to a lack of basic residues, hindering further analysis by tandem MS (Schuller et al., 2016; Suh et al., 2016). Novel proteases, such as chymotrypsin and proteinase K, that cleave at bulky hydrophobic residues like tyrosine have proven effective in the past for analyzing native sequence of CTD but proteolysis become inhibited upon phosphorylation (Mayfield et al., 2017). Short synthetic peptides circumvent the need for proteases but poorly mimic the physiological CTD and are unlikely to reveal bona fide CTD kinase specificities. To overcome these technical challenges, we generated a full-length yeast CTD with Lys replacing Ser7 in every other repeat (yCTD-Lys) (Figure S1C). This allowed for trypsin digestion into di-heptads which represent the functional unit of the CTD (Corden, 2013; Eick and Geyer, 2013), and are amenable to MS/MS analysis.

To validate that the introduction of lysine residues does not bias kinase specificity, we first mapped the phosphorylation pattern of c-Abl or P-TEFb individually along yCTD-Lys using LC-UVPD-MS/MS. When treated with c-Abl, two single phosphorylation species of equal amount are found with tyrosine at the same or neighboring heptad of Lys replacement (Figure 3D and S4A, peak 1 and 2) and a small peak in which both Tyr1 residues are phosphorylated (Figure 3D and S4A, peak 3). When treated with P-TEFb alone, we observed four single phosphorylated peptides: two almost equally abundant peaks containing di-heptads with a single pSer5 (Figure 3E and S4B, peak 1 and 2) and two peaks about ∼40 fold less in intensity with pSer2 or pSer7 (Figure 3E and S4B, peak 3 and 4), consistent with our previously analysis that P-TEFb strongly favors pSer5 in unmodified CTD substrates (Figure 1A). Double phosphorylated species are also detected for di-heptads with both Ser5 residues phosphorylated as the predominant product (Figure 3E and 3F and S4B, peak 5). Several very small peaks (less than 100-fold lower in intensity) are identified as peptides containing both Ser5 and pSer2 (Figure 3E and S4B, peak 6). These results indicate that the existence of Lys residue does not seem to bias kinase activity and is consistent with our previous results that P-TEFb strongly prefers to phosphorylate Ser5.

When treated in tandem with c-Abl followed by P-TEFb and digested with trypsin, diheptads (YSPTSPSYSPTSPK) in a variety of phosphorylation states are generated. These species were separated in CL and revealed 9 di-heptad species of varying abundancesisolated and analyzed. LC purification separates the different phosphorylation states of the di-peptide (Figure 3G). Some of the di-heptides contain only a single phosphorylation due to incomplete reactions *in vitro*. To understand the effect of c-Abl CTD phosphorylation on P-TEFb, we focused on multiply phosphorylated species especially those containing both tyrosine and serine phosphorylation (Figure 3G). Tandem phosphorylation generated species unique to those observed in c-Abl or P-TEFb individual treatment (elution at 27-31 min in LC, Figure 3G, peak 6, 7, 8). The most abundant of these unique species (Figure 3G peak 7) contains both Tyr1 and Ser2 phosphorylation (Figure 3H and S4C). Similarly, a close-by but less abundant peak also contains Tyr1 and Ser2 double phosphorylation although in different location (Figure 3G peak 8 and S4C). Importantly, only a small peak contains both Tyr1 and Ser5 phosphorylation (Figure 3G peak 6 and S4C). Although we cannot exclude the possibility of the existence of other phosphorylated species containing a mixture of tyrosine and serine phosphorylation, their quantity is likely very low and not detected in LC-UVPD-MS/MS analysis. P-TEFb’s serine residue preference is dramatically different from reactions on unmodified CTD substrate, where pSer5 predominates (Figure 3E peak 5), and those pre-treated with c-Abl where pSer2 is the primary product species (Figure 3G peak 7). Our results show that in di-heptads with Tyr1 phosphorylated, Ser2 phosphorylation in the flanking heptad becomes the primary target of P-TEFb phosphorylation.

The high performance of LC chromotography also allowed us to confirm our phosphomapping within in di-heptads in the yeast CTD that diverge from consensus sequence (Figure S1C). Three di-heptads of divergent sequence were generated following trypsin digestion of the yCTD-Lys construct (Figure S5, sequences of YSPTSP<u>A</u>YSPTSPK, YSPTSP<u>N</u>YSPTSPK, and YSPTSP<u>G</u>YSP<u>G</u>SPK). Although these diheptads exist in a much smaller amount than the dominant product YSPTSP<u>S</u>YSPTSPK, they can be resolved and purified in high-performance liquid chromatography and analyzed for phosphorylation position (Figure S5). In these three phosphorylated diheptads, the sole detected product of tandem treatment is a di-heptad with Tyr1 and Ser2 phosphorylated (Figure S5, right panels). Without c-Abl pre-treatment, all peptides phosphorylated by P-TEFb gave a predominant di-heptad species containing only pSer5 (Figure S5, left panels). As in the consensus sequence, no Tyr1 and Ser5 double phosphorylation species were captured, possibly due to low abundance. The phosphoryl mapping of the various di-heptads generated provides independent evidence that Tyr1 phosphorylation promotes Ser2 phosphorylation by P-TEFb even in the context of divergent heptads.

To further corroborate the mass spectrometry results that the specificity of P-TEFb is altered from Ser5 to Ser2 upon Tyr1 phosphorylation, we generated a new yeast CTD variant with 13 repeats (half of the full-length yeast CTD) with every single Ser5 mutated to alanine (S5A construct, sequence in Figure S1H). Previously, it was shown that replacing Ser5 in CTD prevents its phosphorylation by P-TEFb (Czudnochowski et al., 2012). Treatment of the S5A constructs with P-TEFb alone results in the addition of little to no phosphate shown by MALDI-MS (Figures 3I). However, when the S5A construct is treated with c-Abl, it accepts up to 5 phosphate groups (Figure 3J). Subsequent treatment with P-TEFb results in an obvious shift in the MALDI-MS spectra with up to 7 phosphates added to the final product (Figure 3K). The results corroborate the conclusion crawn from the MS/MS results, indicating that upon Tyr1 phosphorylation S5A becomes a viable substrate for P-TEFb and adds at least two additional phosphates likely to Ser2 residues.

The observation of pSer2 as the major product in the context of pre-existing pTyr1 is interesting because P-TEFb has consistently shown a strong preference for Ser5 *in vitro*. Using a combination of immunoblotting, LC-UVPD-MS/MS, mutagenesis, and MALDI-TOF we found Tyr1 phosphorylation primes the CTD for subsequent modification on Ser2 by P-TEFb via alteration of its specificity from Ser5.

### Effect of Ser5 and Ser7 phosphorylation on P-TEFb activity

The observation that Tyr1 phosphorylation by c-Abl alters the specificity of P-TEFb from Ser5 to Ser2 prompted us to ask if other kinases recruited to the CTD at the beginning of transcription can also alter P-TEFb specificity *in vitro*. TFIIH is a kinase during transcription initiation, and its activity promotes P-TEFb function *in vivo* (Ebmeier et al., 2017). To evaluate if a modification or combination of modifications installed by TFIIH can promote the Ser2 specificity of P-TEFb as we see with Tyr1 phosphorylation, we reconstructed CTD phosphorylation *in vitro* by treating yCTD substrates sequentially with TFIIH followed by P-TEFb and analyzed the resultant phosphorylation pattern using LC-UVPD-MS/MS and immunoblotting (Figure S6). When followed by P-TEFb three phosphorylated species are generated, as revealed by LC-UVPD-MS/MS: two major species containing Ser5 phosphorylation and a minor species containing Ser2 phosphorylation (Figure S6A-B). These peptides are reminiscent of those generated by P-TEFb alone where Ser5 phosphorylation dominates (Figure 1A). This data indicates that TFIIH-mediated phosphorylations do not alter P-TEFb specificity *in vitro*.

### Tyr1 phosphorylation primes Ser2 phosphorylation in human cells

Our kinase assays have shown that Tyr1 phosphorylation by c-Abl alters the specificity of P-TEFb allowing for Ser2 phosphorylation of the CTD *in vitro*. To evaluate the importance of pTyr1 to Ser2 phosphorylation in human cells, we sought to selectively reduce pTyr1 levels and monitor pSer2 via western blot (Figure 4A and 4B). Available literature suggests that c-Abl is important to Tyr1 phosphorylation in RNA polymerase II but not the sole kinase responsible (Baskaran et al., 1999). Other Abl-like kinases may likely compensate for the function of c-Abl by phosphorylating Tyr1 in human cells. Therefore, we initially utilized the potent inhibitor dasatinib, which inhibits c-Abl as well as other tyrosine kinases homologous to c-Abl, to treat HEK293T cells (Winter et al., 2012). Tyr1 phosphorylation has also been implicated in stabilizing RNA polymerase II in the cytosol (Hsin et al., 2014), so marked reduction of Tyr1 phosphorylation may lead to a decrease in the global level of RNA polymerase II resulting in an apparent decrease in CTD phosphorylation levels. To address this potential artifact, we optimized inhibitor concentration to a level at which global RNA polymerase II levels are not significantly altered as determined by immunoblotting against RNA polymerase II subunits POLR2A and POLR2C (Figure 4 and S7A). At 10µM dasatinib, pTyr1 levels are reduced by 30% in HEK293T cells and this is accompanied by a 29% decrease in Ser2 phosphorylation (Figure 4A). Importantly, pSer5 levels were not significantly altered (Figure 4A and S7A). This agrees with our *in vitro* observations that pSer5 is independent of pTyr1 status. To more specifically target c-Abl mediated Tyr1 phosphorylation, we utilized the highly specific c-Abl inhibitor imatinib that has a much smaller inhibitory repertoire (Winter et al., 2012). Treatment of HEK293T cells with 20μM imatinib results in a reduction in pTyr1 of 35%. This is accompanied by a statistically significant decrease in pSer2 levels of 15% (Figure 4B). In both the dasatinib and imatinib treatments pSer5, POLR2A, and POLR2C levels remain unaffected (Figure 4 and S7A). Together, this inhibitor-based approach supports our *in vitro* observation that Ser2 phosphorylation is specifically coupled to Tyr1 phosphorylation in cells.

**Figure 4:**
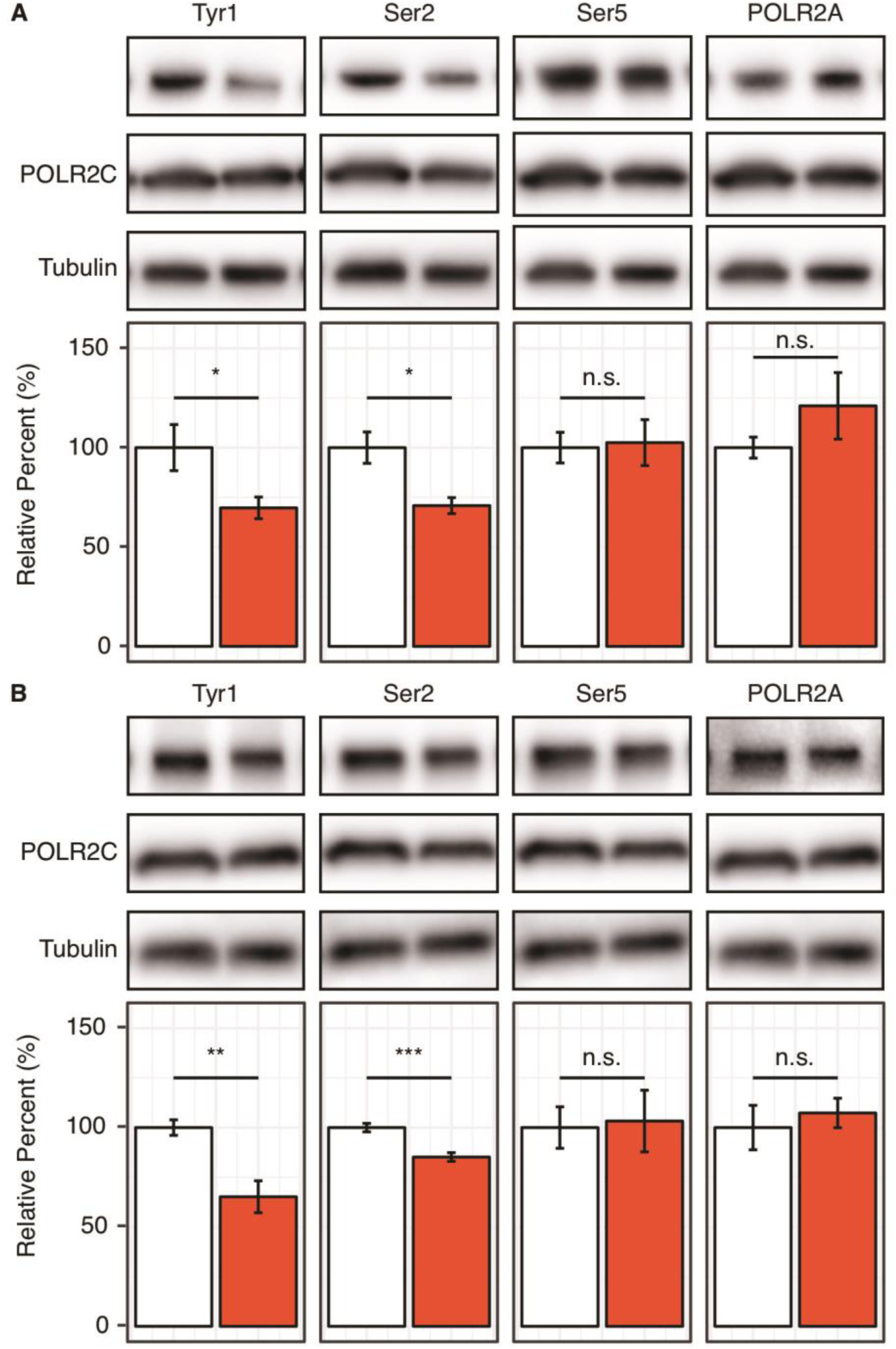
Reduction of Tyr1 levels specifically reduces Ser2 levels in cells. (**A**) Representative images (top) and quantification (bottom) of western blot against 20μg total protein from HEK293T cells treated with paired DMSO vehicle control (left, white) or 10μM dasatinib (right, red). Immuno-blotting against epitopes left to right: phosphorylated Tyr1 reduced by treatment to 69.7% control (n=6), phosphorylated Ser2 reduced by treatment to 70.8% control (n=4), phosphorylated Ser5 unaltered (n=6), POLR2A unaltered (n=6) (POLR2C quantification provided in Supplementary Figure 13). (**B**) Representative images (top) and quantification (bottom) of western blot against 20ug total protein from HEK293T cells treated with paired DMSO vehicle control (left, white) or 20μM imatinib (right, red). Immuno-blotting against epitopes, left to right: Phosphorylated Tyr1 reduced by treatment to 65.2% control (n=6), phosphorylated Ser2 reduced by treatment to 85.2% control (n=6), phosphorylated Ser5 unaltered (n=6), POLR2A unaltered (n=6) (POLR2C quantification supplied in Supplementary Figure 13). Epitope signals normalized against tubulin loading control. Significance determined by Welch’s t-test (* = p-value < 0.05, ** = p-value < 0.01, *** = p-value < 0.001, n.s. = not significant (p-value > 0.05)), error bars indicate SEM.

### Tyr1 phosphorylation promotes Ser2 phosphorylation

To understand the biological implication of coupled Tyr1 and Ser2 phosphorylation at the level of individual genes, we conducted ChIP-seq analysis for the distribution of RNA Polymerase II upon the inhibition of Tyr1 phosphorylation. To carry out this experiment, we inhibited c-Abl with the potent small molecule inhibitor dasatinib in HEK293 cells under conditions where pTyr1 is significantly reduced but overall Pol II amount is un affected (Figure 4A). The sample was prepared for ChIP-seq studies using RNA polymerase II (8WG5) and pSer2 (3E10) specific antibodies for immunoprecipitation to analyze the distribution of RNA polymerase II in a genome-wide fashion. Comparison of the dasatinib treated cells with the vehicle controls, the distribution of RNA polymerase II along the gene body is altered in multiple genes (Figure 5A). Signal was normalized to 10M reads for both samples and a significant increase of peak height for RNA polymerase II was found in the promoter region of many genes, as demonstrated in representative genes Myc and FANCL (Figure 5A). To quantify the change of distribution of the polymerase, we calculated the pausing index (Zeitlinger et al., 2007) which is the ratio of Pol II read density in the region −50 to +300 bp of Transcription starting site (TSS) to the rest of gene body 3000 bp downstream of Transcription end site (TES). The genes are clustered into four groups based on the pausing index: the G0 cluster has a pausing score close to 0; the remaining genes were ranked based on their pausing scores from high to low with the G1 cluster containing genes with pausing scores less than the lower quartile, G2 with genes contained between the lower and upper quartile, and G3 with pausing scores above the upper quartile. Genes in G0 and G1 have little occupancy of the polymerase and might not be active (Figure 5B). Meta-analysis, as visualized in box plot for the pausing index of the genes in G0 and G1, shows no statistical difference between control and treatment samples (Figure S7B and S7D). But a statistically significant increase can be observed in G2 and this increase is close to two-fold in G2 and G3 genes (Figure 5C and S7C-D). The same trend is observed across biological duplicates. Overall, these results indicate that RNA polymerase II is stalled in the promoter region upon the inhibition of Tyr1 phosphorylation.

**Figure 5:**
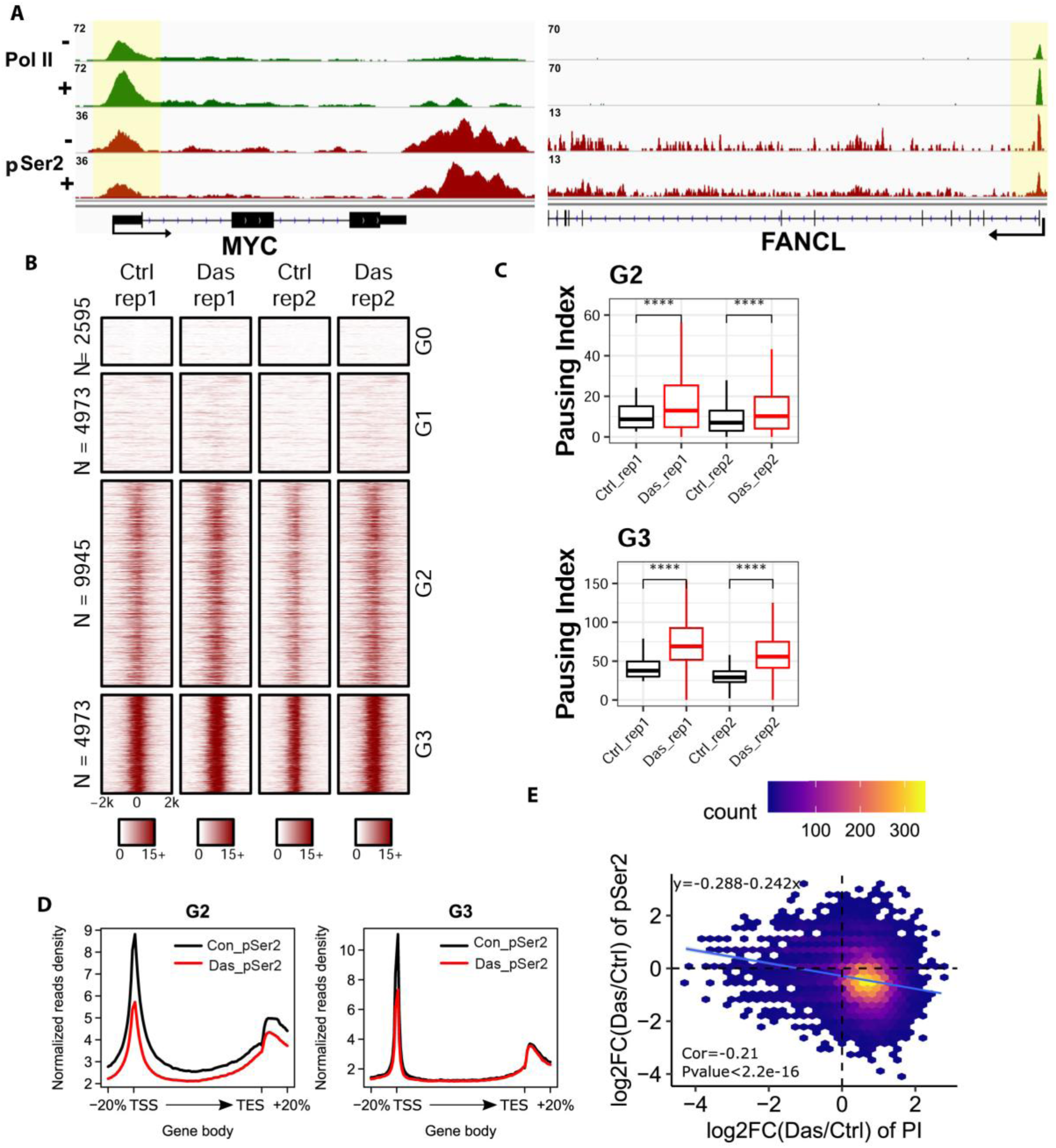
ChIP-seq analyses on the distribution of RNA polymerase II upon the inhibition of Tyr1 phosphorylation. (A) ChIP-seq example illustrating the association of RNA polymerase II along the active transcribing genes. Antibody 8WG5 was used to detect RNA polymerase II regardless of its phosphorylation state and 3E10 was used to detect pSer2. The promoter regions of the genes are shaded in yellow for highlighting. (B) Heatmaps of ChIP-seq signal intensity of RNA polymerase II (8WG5) [±2kb windows around the center of transcription start site (TSS)] for genes in each group. (C) Boxplots on the pausing index changes on the genes from G2 (9945 genes) and G3 (4973 genes) clusters upon Tyr1 phosphorylation. “****” indicates p-value ≤ 0.0001. (D) The profiling of pSer2 signal along gene body under control and dasatinib treatment for genes of cluster G2 and G3. (E) The correlation of the accumulation of RNA polymerase II versus the reduction of pSer2 in the promoter region for genes in G2 and G3. The x-axis represents the log2 of the ratio of pausing index in the dasatinib treatment versus DMSO control. The y-axis presents the log2 ratio of pSer2 signal strength in dasatinib treatment versus DMSO control.

Upon inhibition of Tyr1 phosphorylation, overall pSer2 level is also reduced (Figure 4). To further define the stalling of RNA polymerase II in promoter region, we analyzed the distribution of pSer2 along the gene bodies upon inhibition of tyrosine kinase using dasatinib (Figure 5D). Across genes, there is a strong reduction of pSer2 signals around the promoter regions of the genes while a relatively weak reduction was observed in gene body regions (as representative genes Myc and FANCL in Figure 5A). This trend is in particularly obvious in genes clustered in G2 and G3 based on their pausing index (Figure 5D). Upon comparison, most of the genes with increased pausing index also show reduced pSer2 signal intensity around promoter regions (Figure 5D and Figure S5E). The change of pausing index significantly negatively correlates with the change of pSer2 signals upon inhibition of Tyr1 phosphorylation (Figure 5E). Thus, these results suggested that inhibition of Tyr1 phosphorylation result in reduced pSer2 levels and impede the release of paused Pol II into the productive elongation.

## Discussion

Our discovery that Tyr1 phosphorylation of the CTD alters the preference of P-TEFb from Ser5 to Ser2 resolves controversy surrounding P-TEFb’s specificity (Bartkowiak et al., 2010; Czudnochowski et al., 2012). P-TEFb was initially identified as a CTD kinase that controls the elongation potential of RNA polymerase II, is required for the majority of RNA polymerase II transcription, and is specific for Ser2 *in vivo* (Chao and Price, 2001; Marshall et al., 1996; Ni et al., 2004). However, these early conclusions are at odds with *in vitro* data demonstrating P-TEFb is incapable of phosphorylating Ser2 of CTD peptides *in vitro* (Czudnochowski et al., 2012). Two other kinases, CDK12 and CDK13, display Ser2 kinase activity in cells but do not seem to play a major role in Ser2 phosphorylation in early transcriptional events (Bartkowiak et al., 2010; Chen et al., 2007). Investigations on the effect of CTD phosphorylations on P-TEFb specificity have revealed that Ser7 (Czudnochowski et al., 2012) and Ser5 (this manuscript) do not alter its preference to Ser5. Using direct methods, like mass spectrometry confirmed by immunoblotting and EMSA, we identified that Tyr1 phosphorylation could alter the specificity of P-TEFb from Ser5 to Ser2 *in vitro*. Furthermore, inhibition of Tyr1 phosphorylation leads to the reduction of Ser2 phosphorylation in human cells and the accumulation of Pol II in the promoter-proximal pausing stage. Therefore, we show that Tyr1 phosphorylation potentiates Ser2 phosphorylation of the CTD by altering P-TEFb specificity.

The PTM state of the CTD has been correlated to the progression of transcription (Jeronimo et al., 2016). Traditionally, such analyses are interpreted through a paradigm considering heptads phosphorylated on a single isolated residue with Ser5 phosphorylation dominating the initiation stage of transcription and Ser2 phosphorylation dominating elongation and termination (Corden, 2013). However, this over-simplification of CTD modification cannot explain the well-coordinated recruitment of the myriad CTD binding factors currently implicated in eukaryotic transcription (Ebmeier et al., 2017; Eick and Geyer, 2013; Harlen and Churchman, 2017). Data presented here points to a sophisticated model in which the phosphorylation of Tyr1 at the beginning of transcription sets the stage for future coding events. This interplay between c-Abl and P-TEFb results in a chemically distinct phospho-CTD landscape compared to CTD phosphorylated by a single kinase. The combination of these modification modes likely contributes to a heterogeneous collection of modified heptads, which recruits the diverse array of CTD binding partners in a coordinated manner. These results are in good agreement with the “CTD code” hypothesis proposed decades ago where the different combination of post-translational events result in different transcriptional outcomes.

Tyr1 phosphorylation has been implicated in stabilizing RNA polymerase II in cells (Hsin et al., 2014), transcription termination (Mayer et al., 2012) and anti-sense transcription (Descostes et al., 2014) but a coherent molecular basis for these disparate functions remains elusive. Our analysis provides a molecular mechanism demonstrating how Tyr1 phosphorylation affects subsequent phosphorylation events carried out by other CTD kinases. Our ChIP-seq analysis reveals that Tyr1 phosphorylation changes will alter the overall distribution of RNA polymerase II with more polymerase stalled to the promoter proximal pausing regions of genes and a reduction of pSer2 levels at these accumulation sites. This directly implicates Tyr1 phosphorylation in regulating pausing via pSer2 phosphorylation. The ability of Tyr1 phosphorylation to redirect signaling and influence subsequent modifications along the CTD, as revealed for P-TEFb, suggests these various roles for pTyr1 may arise indirectly through its impact on downstream CTD modifiers, highlighting integrated, indirect, and context-specific mechanisms for pTyr1 during co-transcriptional signaling that diversify CTD phosphorylation and facilitate transcriptional and allied processes. The abundance of pTyr1 appears to be far less than pSer2 or pSer5. However, the final accumulation of individual species is dependent on the dynamic interplay of CTD kinases and phosphatases throughout the transcription cycle. Tyr1 phosphorylation is relatively transient, appearing at the transition from initiation to elongation and decreasing rapidly through the action of phosphatase(s) (Eick and Geyer, 2013). Despite this transient nature, pTyr1 is positioned in a vital window to alter P-TEFb specificity and regulate its phosphorylation pattern along RNA polymerase II. The adjustability of P-TEFb specificity by nearby Tyr1 phosphorylation states also reveals a novel mechanism for the regulation of P-TEFb kinase activity. With many binding partners in cells for P-TEFb, there might be additional regulators promoting the pSer2 activity of P-TEFb independent of or cooperatively with Tyr1 phosphorylation.

The data presented reconciles P-TEFb’s *in vitro* and *in vivo* specificity and inspires new queries fundamental to CTD biology. P-TEFb is ubiquitously important for transcription across eukaryotic cells and often co-opted in disease states like HIV infection and cancer (Franco et al., 2018). The integrated CTD code revealed here represents a unique mechanism to manipulate P-TEFb and potentially other CTD modifiers. Overall, our findings support a model in which cross-talk between CTD modification enzymes increases the diversity and coding potential of CTD heptads. This expands the lexicon of phosphorylation marks and can provide more specific recruitment of transcription regulators allowing for the precise control of eukaryotic transcription.

### Experimental methods

*E. coli* BL21 (DE3) cells grown 37°C in Luria-Bertani media were used to overexpress recombinant GST-CTDs and Erk2 variants. HEK 293T cells were maintained in DMEM (ISC BioExpress cat#T-2989-6) supplemented with 10% FBS, penicillin and streptomycin. The cells were split every other day when it reached confluency of 70-80%. Seeding at a concentration of 9.6 × 10^4^ cells per 10 cm culture dish and incubated at 37°C at 5% CO_2_.

### Protein expression and purification

CTD coding sequences (Figure S1) were subcloned into pET28a (Novagene) derivative vectors encoding an N-terminal His-tag a GST-tag and a 3C-protease site to generate GST-CTD constructs as described previously (Mayfield et al., 2017). 3CTD-5CTD and YtoF variants of CTD coding portions were amplified from synthetic DNA templates generated by IDT. The S5A variant DNA and the S7K spaced DNA constructs were purchased from Biomatik as synthetic genes, were amplified and subsequently cloned into the pET28a derivative vector described above. *Homo sapiens* Erk2 was expressed from pET-His6-ERK2-MEK1_R4F_coexpression vector. pET-His6-ERK2-MEK1_R4F_coexpression was a gift from Melanie Cobb (Addgene plasmid #39212) (Khokhlatchev et al., 1997).

GST-CTD variants were prepared by established protocols. Briefly, proteins were overexpressed in *E. coli* BL21 (DE3) cells by growing at 37°C in Luria-Bertani media containing 50 μg/mL kanamycin or ampicillin to an OD_600_ of 0.4-0.6. Expression was induced by the addition of isopropyl-β-D-thiogalactopyranoside (IPTG) to a final concentration of 0.5 mM. After induction, the cultures were grown at 16°C for an additional 16 hours. The cells were pelleted and lysed via sonication in lysis buffer [50 mM Tris-HCl pH 8.0, 500 mM NaCl, 15 mM Imidazole, 10% Glycerol, 0.1% Triton X-100, 10 mM β-mercaptoethanol (BME)]. The lysate was cleared by centrifugation at 15,000rpm for 45 minutes at 4°C. The supernatant was initially purified using Ni-NTA (Qiagen) beads and eluted with elution buffer (50 mM Tris-HCl pH 8.0, 500 mM NaCl, 200 mM Imidazole, and 10 mM BME). The protein was dialyzed against gel filtration buffer (20 mM Tris-HCl pH 8.0, 50 mM NaCl, 10 mM BME for GST-yCTD & 20mM Tris-HCl pH 7.5, 200mM NaCl, 10mM BME). Finally, proteins were concentrated and ran on a Superdex 200 gel filtration column (GE). Erk2 was purified using a previous published protocol (Khokhlatchev et al., 1997). Homogeneity of the eluted fractions was determined via Coomassie Brilliant Blue stained SDS-PAGE. Samples were concentrated in vivaspin columns (Sartorius).

### Kinase reactions

Abl kinase treated CTD reactions were prepared in buffer conditions containing 1 μg/μL GST-yCTD substrate, 0.0035 μg/μL c-Abl kinase, 50 mM Tris-HCl at pH7.5, 50 mM MgCl_2_ and 2 mM ATP. TFIIH treated CTD reaction were prepared in buffer conditions containing 1 μg/μL GST-CTD substrate, 0.025 μg/μL TFIIH, 50 mM Tris-HCl at pH7.5, 50 mM MgCl_2_ and 2 mM ATP. P-TEFb treated CTD reaction, were prepared in buffer conditions containing 1 μg/μL GST-CTD substrate, 0.0075 μg/μL P-TEFb, 50 mM Tris-HCl at pH7.5, 50 mM MgCl_2_ and 2 mM ATP. Erk2 treated CTD reaction, as well as the controls with no kinase treatment, were prepared in buffer conditions containing 1 μg/μL GST-CTD substrate, 0.025 μg/μL Erk2, 50 mM Tris-HCl at pH7.5, 50 mM MgCl_2_ and 2 mM ATP. These reactions were incubated for various amount of time at 30°C along with control experiments setup under identical conditions but without kinases and then stored at −80°C until analysis.

Tandem kinase treatments were performed by mixing 10 μg GST-CTD substrate of c-Abl in a buffer containing 0.0035 μg/μL c-Abl kinase, 50mM Tris-HCl at pH7.5, 50 mM MgCl_2_, 4 mM ATP for overnight and then added the second kinases (0.05 μg/μL TFIIH or 0.015 μg/μL P-TEFb or 0.05 μg/μL Erk2). These were incubated at 30°C for 16 hours and stored at −80°C until analysis. For c-Abl inactivation assays, 10 minutes of incubation of reaction at 80 ͦC or 10mM of c-Abl inhibitor were added before the addition of P-TEFb.

### Kinase activity assay

The Erk2 kinetic activity assay was performed in a 25µl reaction volume containing 0-100µM substrate (GST yCTD or GST YtoF CTD) and a reaction buffer of 40mM Tris-HCl at pH 8.0 and 20mM MgCl_2_. The reaction was initiated by adding 187nM of Erk2 and incubated at 28°C for 15mins before being quenched with 25µl H_2_O and 50µl of room temperature Kinase-Glo Detection Reaction (Promega). The mixtures were allowed to sit at room temperature for 10 minutes before reading the bioluminescence in a Tecan Plate reader 200. The readings obtained were translated to ATP concentration with the help of an ATP standard curve determined with the Kinase Detection Reagent.

The TFIIH kinetic reactions were set up with 0-100µM substrate (GST yCTD or GST YtoF CTD), 0.2µM TFIIH, 0.1mg/ml BSA and reaction buffer of 50mM Tris-HCl pH 8.0, 10mM MgCl_2_, 1mM DTT. 500µM ATP Mix (10 nCi/µl radiolabeled ATP, PerkinElmer) was added to each tube to start the reactions. The tubes were subsequently incubated in a 30 °C water bath for 30mins and quenched with 500µl of quench buffer (1mM potassium phosphate pH 6.8, 1mM EDTA) to a reaction volume of 10µl. Each reaction was loaded onto 0.45 µm nitrocellulose filters and washed three times with 1mM potassium phosphate buffer to remove any excess labeled ATP. Filters were added to glass vials with scintillation fluid, Econo-Safe Economical Biodegradable Counting Cocktail (Research Products International) and set in a scintillation counter for 5min reads each. The amount of phosphate incorporation was determined for each reaction using a set of 147pmol labeled ATP standards that were read alongside each reaction set.

Kinetic data obtained from the two assays described above were analyzed in R (Hamilton, 2015; Team, 2017) and fitted to the Michaelis-Menten kinetic equation to obtain respective kinetic parameters *k_cat_*(s^-1^) and *K_m_* (µM).

### Electrophoretic mobility shift assay

SDS-PAGE analysis was performed using 10-15% acrylamide gels containing 1% SDS. GST-CTD samples were prepared by boiling with SDS-PAGE loading dye at 95°C for 5 minutes. This was also used to quench time-course reactions. A volume containing approximately 1μg of phosphorylated GST-CTD substrate or no kinase control was loaded into wells and resolved at ∼150V for 1h at room temperature. Gels were stained with Coomassie Brilliant Blue and visualized on G: BOX imaging systems (Syngene).

### MALDI-MS analysis

Approximately 5 μg of GST-CTD protein from the kinase reactions described were prepared for MALDI-MS. If necessary, the protein was digested with 3C protease by mixing sample in a 1:10 ratio of 3C-protease to GST-CTD construct. Proteins were equilibrated with dilute trifluoracetic acid (TFA) to a final concentration of 0.1% TFA and a pH of < 4. These samples were desalted using ZipTip (Millipore) tips according to manufacturer instructions. These samples were mixed 1:1 with a 2,5-dihydroxybenzoic acid matrix solution (DHB) and spotted on a stainless steel sample plate. The spots were allowed to crystallize at ambient temperature and pressure. MALDI-MS spectra were obtained on an AB Voyager-DE PRO MALDI-TOF instrument with manual adjustment of instrument parameters to ensure the greatest signal to noise. Sample masses were determined by a single point calibration against the untreated GST-CTD construct. Data analysis, noise reduction, and Gaussian smoothing, if necessary, were performed in DataExplorer (AB), R, and the R package smoother to provide interpretable data (Hamilton, 2015; Team, 2017). Data was visualized in R-Studio using ggplot2 (Wickhan, 2009). Masses were determined as the highest local intensity peak of the post-processed data.

### Cell culture and total protein preparation

HEK293T cells were maintained in DMEM (ISC BioExpress cat#T-2989-6) with splitting every other day and seeding at a concentration of 9.6 × 10^4^ cells per 10 cm culture dish and incubated at 37°C at 5% CO_2_. Cells to be treated with imatinib, dasatinib, or vehicle control were platted at 5 × 10^5^ cells per well in 6-well tissue culture plates in fresh DMEM. Cells were incubated for 24 hours and media was replaced with fresh DMEM containing the indicated amount of inhibitor or equivalent portion of DMSO vehicle control for an additional 24-40 hours.

Protein preparations were generated by direct in-well lysis. Media was then removed, and the cells were washed with ice-cold PBS, and 200 μL RIPA buffer (150 mM NaCl, 10mM Tris-HCl pH 7.5, 0.1% SDS, 1% Triton X-100, 1% deoxycholate, and 5 mM EDTA) supplemented to 1X with HALT protease and phosphatase inhibitor cocktail (Thermo Scientific) was added directly to cells. Plates were incubated on ice for 15 minutes with gently shaking, and the lysate was transferred to micro centrifuge tubes. Samples were briefly sonicated to reduce viscosity and spun at 13,000 rpm for 15 minutes to remove cell debris. Protein concentration was determined to utilize Pierce^TM^ BCA Protein Assay Kit (Thermo Scientific) against a BSA standard curve. Samples were diluted with SDS-PAGE loading buffer and boiled at 95°C for 5 minutes. The sample was aliquoted and frozen at −80°C.

### Immunoblotting

Total protein from cell lysate (20-40 μg) or GST-yCTD samples (50 to 500ng, dependent on epitope) was loaded onto a 4-20% gradient SDS-PAGE gel (Biorad, Cat#:456-1096) and ran at 150V for 50 minutes at room temperature in a Mini-PROTEAN Tetra Cell (Biorad). The proteins were transferred to PVDF membrane at 100 V for 1 hour on ice in a Mini-PROTEAN Tetra Cell (Biorad). Membranes were blocked in 1X TBST (20 mM Tris-HCl pH 7.5, 150 mM NaCl, 0.1% Tween-20) and 5% Bovine Serum Albumin (BSA) or non-fat dry milk (NFDM) for 1 hour at room temperature with shaking. Blocked membranes were incubated in primary antibody in either 1X TBST or 1X TBST+5% BSA at 4 °C overnight or 1 h at room temperature. The membranes were then washed six times with 1X TBST for 5 minutes each at room temperature and incubated with secondary antibody in 1X TBST for 1 hour at room temperature. The membrane was washed once again and incubated with SuperSignal West Pico Chemiluminescent Substrate (Pierce, Cat#: 34079) according to factory directions. Blots were imaged using a G: BOX gel doc system (Syngene) and quantified in ImageJ (Schneider et al., 2012). Statistical analysis was performed in R (Team, 2017).

Blots normalized against Coomassie-stained bands were stained by incubating the membrane post-immunoblotting with stain solution (0.1% Coomassie Brilliant Blue R-250, 40% ethanol, 10% acetic acid) for 1 minute. The stain was discarded, and the blot was briefly rinsed with distilled water. Blot was de-stained in de-stain solution (10% ethanol, 7.5% acetic acid) until bands were visible. Blots were equilibrated with distilled water and imaged wet in a plastic blot protector using a G: BOX gel doc system (Syngene) and quantified as above.

For dot blot, samples of GST-yCTD were treated with P-TEFb alone or c-Abl followed by P-TEFb. The heat inactivated samples were prepared by heating the c-Abl treated GST-yCTD at 60°C for five minutes, while the Dasatinib inactivated samples were prepared by adding 10µM Dasatinib to the c-Abl treated sample 15 minutes before incubation with P-TEFb. The dot blots were performed by adding 2X SDS page loading dye and briefly heating each sample after which 1µg of the sample was loaded in three replicates onto a 0.45µm Nitrocellulose membrane. The membrane was subsequently allowed to dry and blocked in 1X TBST (20 mM Tris-HCl pH 7.5, 150 mM NaCl, 0.1% Tween-20) and 5% Bovine Serum Albumin (BSA) for 1 hour at room temperature with shaking. The Anti-RNA Pol II phosphoSer2 antibody (clone: 3E10, Millipore Cat: 14-1571) was diluted 1:5000 times and was incubated overnight at 6°C to probe for Ser2 phosphorylation. The membranes were then washed five times with 1X TBST for 5 minutes each at room temperature and incubated with secondary antibody in 1X TBST for 1 hour at room temperature. The membrane was washed once again and incubated with SuperSignal West Pico Chemiluminescent Substrate (Pierce, Cat#: 34079) according to manufacturer instruction. Blots were imaged using a G: BOX gel doc system (Syngene) and quantified in ImageJ (Schneider et al., 2012). Statistical analysis was performed by using the Data Analysis function in Microsoft Excel.

### LC-UVPD-MS/MS Analysis

GST-3CTD samples (approximately 1 µg/µL) were digested on ice for 4 hours in 50 mM Tris-HCl at pH 8.0 with 150 mM NaCl using 3C-protease at a molar ratio of 100:1 protein: protease in a reaction volume of 20 µL. Digests were desalted on C18 spin columns and resuspended to 1 µM with 0.1% formic acid for LC-MS analysis.

Separations were carried out on a Dionex Ultimate 3000 nano liquid chromatograph plumbed for direct injection. Picofrit 75 µm id analytical columns (New Objective, Woburn, MA) were packed to 20 cm using 1.8 µm Waters Xbridge BEH C18 (Milford, MA). Mobile phase A was water and B was acetonitrile, each containing 0.1% formic acid. Separations occurred over a 30-minute linear gradient from 2-35% B. The flow rate was maintained at 0.3 µL/min during the separation.

An Orbitrap^TM^ Fusion Lumos Tribrid mass spectrometer (Thermo Fischer Scientific, San Jose, CA) equipped with a Coherent ExciStar XS excimer laser operated at 193 nm was used for positive mode LC-MS/MS analysis of the 3CTD peptides. The Lumos mass spectrometer was modified for ultraviolet photodissociation (UVPD) as described earlier (Klein et al., 2016). Photoactivation in the low-pressure linear ion trap was achieved using 2 pulses at 2 mJ in a targeted *m/z* mode. The 3+ charge states of the singly and doubly phosphorylated peptide GPGSGMYSPTSPSYSPTSPSYSPTSPS were targeted for photoactivation. All data were acquired in the Orbitrap analyzer where MS1 and MS/MS spectra were collected at resolving powers of 60K and 15K (at *m/z* 200), respectively. MS1 spectra were acquired from *m/z* 400-2000 with an AGC setting of 5E5. Each MS/MS spectrum consisted of two microscans collected from *m/z* 220-2000 with an AGC setting of 2E5.

Data analysis was performed using the XCalibur Qual Browser and ProSight Lite (Fellers et al., 2015). For both targeted *m/z* values, the MS/MS spectrum for each phosphoform present was deconvoluted to neutral forms using Xtract with a signal-to-noise threshold of 3. Sequence coverage was determined by matching the nine ion types observed with UVPD (*a*, *a^•^*, *b*, *c*, *x*, *x^•^*, *y*, *y*-1, *z*). Localization of the phosphorylation(s) was performed by adding a phosphate group (+79.966 Da) at each of the possible serine, threonine, and tyrosine residues to identify fragment ions containing the moiety and optimize characterization scores in ProSight Lite.

Analysis of yCTD treated with P-TEFb was performed identically to previous analysis of yCTD treated with TFIIH and Erk2 (Mayfield et al., 2017). GST-yCTD samples were prepared for bottom-up analysis using a two-step proteolysis method. First, overnight digestion with trypsin at 37°C was carried out using a 1:50 enzyme to substrate ration, which cleaved the GST-portion of the protein while leaving the abasic 26mer CTD portion intact. The resulting digest was filtered through a 10 kDa molecular weight cutoff (MWCO) filter to remove tryptic GST peptides and buffer exchange the CTD portion into 50mM Tris-HCl pH 8.0 and 10mM CaCl_2_ for subsequent proteainase K digestion. Proteinase K was added in a 1:100 ration and digested overnight at 37°C. Samples were diluted to 1μM in 0.2% formic acid for LC-MS. Analysis of yCTD-Lys treated by c-Abl, P-TEFb or c-Abl followed by P-TEFb are using a similar method as described above except the first digestion was done by 3C-protease and second by trypsin.

Bottom-up analysis of yCTD was performed on a Velos Pro dual linear ion trap mass spectrometer (Thermo Fisher, San Jose, CA) equipped with a Coherent ExciStar XS excimer laser (Santa Clara, CA) at 193 nm and 500 Hz as previously described for UVPD (Gardner et al., 2008; Madsen et al., 2010). Two pulses of 2mJ were used for photodissociation. Separations were carried out on a Dionex Ultimate 3000 nano liquid chromatograph configured for preconcentration. Integrafrit trap columns were packed to 3.5 cm using 5 μm Michrom Magic C18. Picrofrit analytical columns were packed to 20 cm using 3.5 μm Waters Xbridge BEH C18 (Milford, MA). Mobile phase A was water and mobile phase B was acetonitrile; each contained 0.1% formic acid. Peptides were loaded onto the trap column for 5 minutes in an aqueous solvent containing 2% acetonitrile and 0.1% formic acid at a 5 μL/min flow rate. Separations occurred over a 20-minute linear gradient in which percent phase B was increased from 2-15% during the first 15 minutes and further increased to 35% over the last 5 minutes. The flow rate was constant at 0.3 μL/min. A top seven data-dependent acquisition method was first used to identify the main phosphorylated species. A targeted analysis followed in which the singly phosphorylated heptad peptides were continually selected for UVPD activation (between MS^1^ acquisitions occurring after every five MS/MS events) to resolve partially co-eluting phospho-isomers. Resulting UVPD spectra were manually interpreted.

### ChIP-seq analysis

For ChIP-seq experiments, HEK293T cells were seeded at 3.5 million cells in a 15cm dish. After 24hours, when the cells achieved confluency of 40-50%, the media was replaced by fresh media containing 10µM Dasatinib inhibitor or the DMSO control and allowed to grow for another 24 hours until confluency of 80% was achieved. The cells were fixed with 1% formaldehyde in 15ml of media, for 8 minutes at room temperature with intermittent swirling. The reaction was quenched by the addition of glycine to a final concentration of 0.125M and incubation for five minutes at room temperature. The cells were washed twice with 15ml of ice-cold Dulbecco’s phosphate-buffered saline (DPBS) and scrapped off the surface with 900µl of DPBS into microcentrifuge tubes. The cells were pelleted at a speed of 8000xg for 5minutes, resuspended and aliquoted such that the number of control cells (with only DMSO) were normalized to the number of dasatinib treated cells, in snap cap Eppendorf tubes. The cell pellet was frozen in a freezing mixture comprised of dry ice and ethanol.

The cells were lysed by adding buffer LB1 [50mM HEPES at pH 7.5, 140mM NaCl, 1mM EDTA, 10% glycerol, 0.5% NP-40, 0.25% Triton-X 100, 1x Protease inhibitor cocktail (thermoscientific)] and placing the tubes on a rotating wheel at 4°C for 10 minutes, following which they were spun at 2000g for 5 minutes to isolate the nuclei as a pellet. These were washed with buffer LB2 [10mM Tris-HCL at pH 8.0, 200mM NaCl, 1mM EDTA, 0.5mM EGTA + 1X Protease inhibitor cocktail (thermoscientific)], and subsequently the nuclei where resuspended in 300µl of nuclear lysis buffer LB3 [10mM Tris-HCL, pH 8, 100mMNaCl, 1mM EDTA, 0.5mM EGTA, 0.1% Na-Deoxycholate, 0.5% N-lauroylsarcosine and 1x Protease inhibitor cocktail (thermoscientific)].

The nuclear lysate of 300µl was sonicated using a Biorupter UCD 200 for 25 cycles at maximum intensity (15sec ON 45 sec OFF in a water bath at 4°C). After each of the 10 cycles, the samples were incubated on ice for 10 minutes. Following sonication 30µl of buffer LB3 supplemented with 10% TritonX-100 was added into the sample and spun at full speed for 10 minutes to remove cell debris. 30µl of the supernatant was taken as the input control for ChIP-seq, and the rest is used to prepare the samples.

Magnetic Protein-G beads were incubated with respective antibody (1µg per 10µl of beads) overnight on the rotating shaker at 4°C. The beads were then washed thrice with 5% BSA in PBS to remove any excess antibody, and the 300µl of the sonicated lysate prepared above is added to it, with 800µl of buffer LB3 and 100µl of buffer LB3 supplemented with10% Triton X-100. The samples were placed on a rotating wheel overnight at 4°C for the immunoprecipitation to occur. The beads were washed twice by a low salt buffer (0.1% Na Deoxycholate, 1% Triton X-100, 1mM EDTA, 50mM HEPES at pH 7.5, 150mM NaCl) followed by wash with high salt buffer (0.1% Na Deoxycholate, 1% Triton X-100, 1mM EDTA, 50mM HEPES at pH 7.5, 500mM NaCl), lithium chloride buffer (250mM LiCl, 0.5% NP-40, 0.5% Na Deoxycholate, 1mM EDTA, 10mM TrisCl at pH 8.1) and finally washed twice with TE buffer (10mM TrisCl at pH 8.1 and 1mM EDTA). The beads were ultimately resuspended in 200µl of elution buffer (1% SDS and 0.1M sodium bicarbonate) and placed in the thermomixer at 65°C for 16 hours to enable reverse crosslinking.

Both the input and treatment samples were with 70µl of elution buffer (1% SDS and 0.1M sodium bicarbonate) and underwent reverse crosslinking at 65°C for 16 hours. After the reverse crosslinking, phenol-chloroform extraction was used to extract the immunoprecipitated DNA, Library prep was done using a starting amount of 3ng of DNA (measured by Qubit HS) using the NEBNext Ultra II DNA Library Prep Kit for illumine (NEB E#7103) following the vendor manual. The libraries with multiplex index primers prepared above were pooled together and sequenced using the NextSeq single end 75 base pair sequencing platform.

Reads were aligned to the human genome (hg19) using bowtie with “--best -- strata –m 1” parameters (Langmead et al., 2009). Only uniquely mapped reads were selected for downstream analysis. MACS2 was employed to call peaks by comparing immunoprecipitated chromatin with input chromatin using standard parameters and a q-value cutoff of 1e-5 (Zhang et al., 2008). The peaks overlapped with the blacklist regions downloaded from UCSC were removed. Each sample was normalized to 10 million mapped reads and visualized in Integrative Genomics Viewer (IGV) (Robinson et al., 2011). The pausing index was defined as the ratio of Pol II density in the promoter proximal region and the Pol II density in the transcribed region (Zeitlinger et al., 2007). The proximal promoter region is defined as −50bp to +300bp around the transcription start site (TSS); while the transcribed region (gene body) is from +300bp to the 3000bp downstream of transcription end site (TES) (Rahl et al., 2010).

### Quantification and data analysis

MALDI data analysis, noise reduction, and Gaussian smoothing (if necessary) were performed in DataExplorer (AB), R, and the R package smoother to provide interpretable data (Hamilton, 2015; Team, 2017). Data was visualized in R-Studio using ggplot2 (Wickhan, 2009). Masses were determined as the highest local intensity peak of the post-processed data. Tandem mass spectrometry data analysis was performed using the XCalibur Qual Browser and ProSight Lite. For both targeted *m/z* values, the MS/MS spectrum for each phosphoform present was deconvoluted to neutral forms using Xtract with a signal-to-noise threshold of 3. Sequence coverage was determined by matching the nine ion types observed with UVPD (*a*, *a^•^*, *b*, *c*, *x*, *x^•^*, *y*, *y*-1, *z*). Localization of the phosphorylation(s) was performed by adding a phosphate group (+79.966 Da) at each of the possible serine, threonine, and tyrosine residues to identify fragment ions containing the moiety and optimize characterization scores in ProSight Lite (Fellers et al., 2015). Western blots were quantified using ImageJ (Schneider et al., 2012) and statistical significance was determined by two-tailed unpaired Student’s t-test assuming unequal variances in Microsoft Excel. Statistical significance is reported in the figure legends. Results have been shown with ± standard deviation or SEM as mentioned in the figure legends.

## Acknowledgments

We want to thank Drs. G. Gill and J. Dixon for comments on the manuscript. This work is supported by grants from the National Institutes of Health (R01 GM104896 and 125882 to Y.J.Z. and R21EB018391 to J.S.B.) and Welch Foundation (F-1778 to Y.J.Z. and F-1155 to J.S.B.). Voyager DE-PROT MALDI MS data were collected in the University of Texas at Austin Proteomics Facility. Funding from the UT System for support of the UT System Proteomics Core Facility Network is gratefully acknowledged.

## Author Contributions

Y.J. Zhang, J.E. Mayfield and J. S. Brodbelt designed the study. E. Mayfield conducted the biochemical, biophysical, and western blot experiments. S. Irani and J. E. Mayfield conducted MALDI-TOF analysis, S. Irani prepared ChIP-seq libraries, Z. Zhang and Z. Liu analyzed the ChIP-seq data, E. E. Escobar, M. R. Robinson, M. R. Mehaffey and S. Sipe did the LC-UVPD-MS/MS analyses. N.T. Burkholder and W. Yang conducted the immunoblotting and kinetic assays. The manuscript was written by J. E. Mayfield and Y.J. Zhang with input from all contributing authors.

---

**Supplementary Figure S1.**
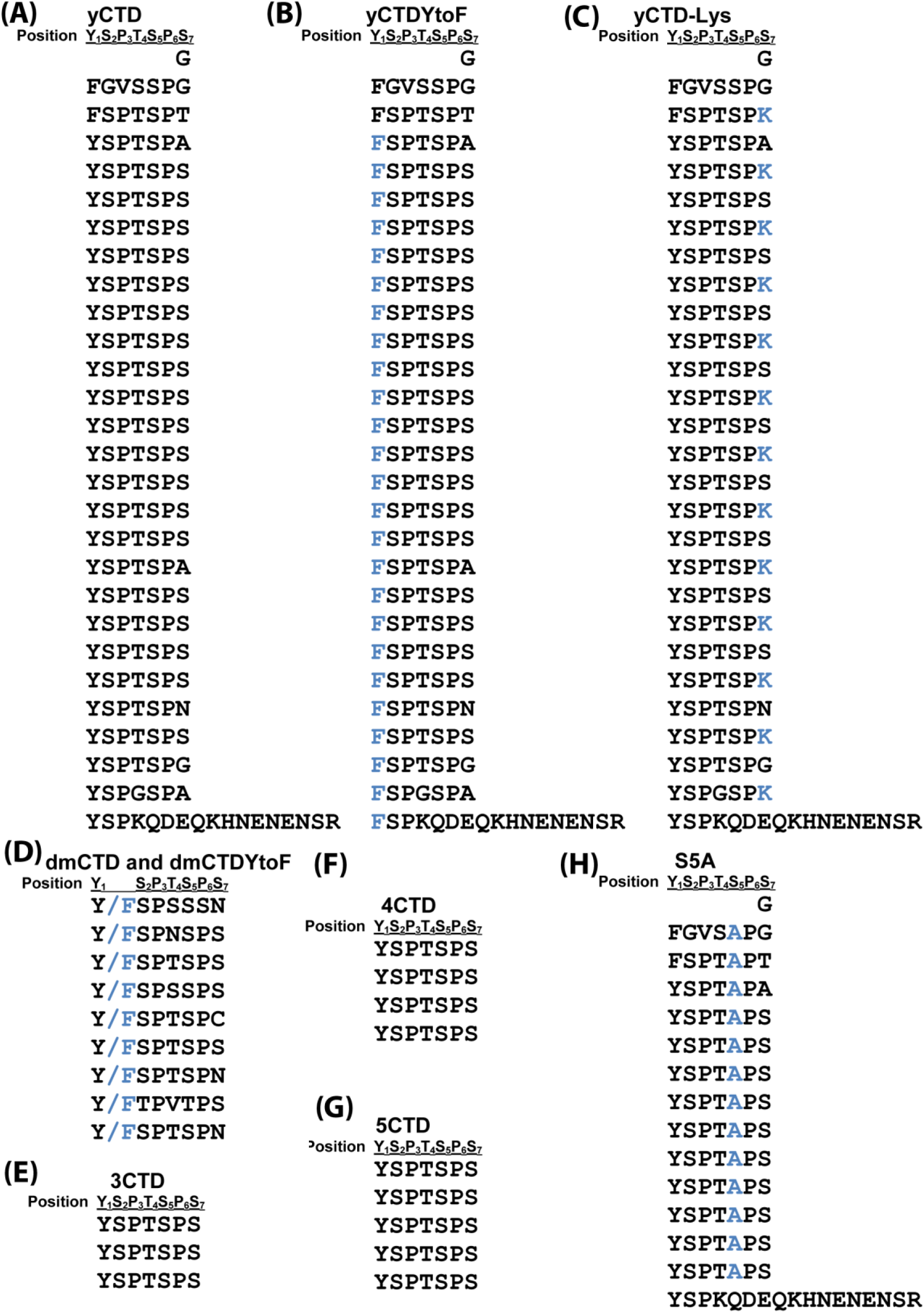
Amino acid sequences of GST-CTD constructs. (A) Sequence of CTD portion of GST-CTD constructs used in the manuscript. Sequences presented N to C-terminal as individual heptads. Residues mutated are indicated above in blue font.

**Supplementary Figure S2.**
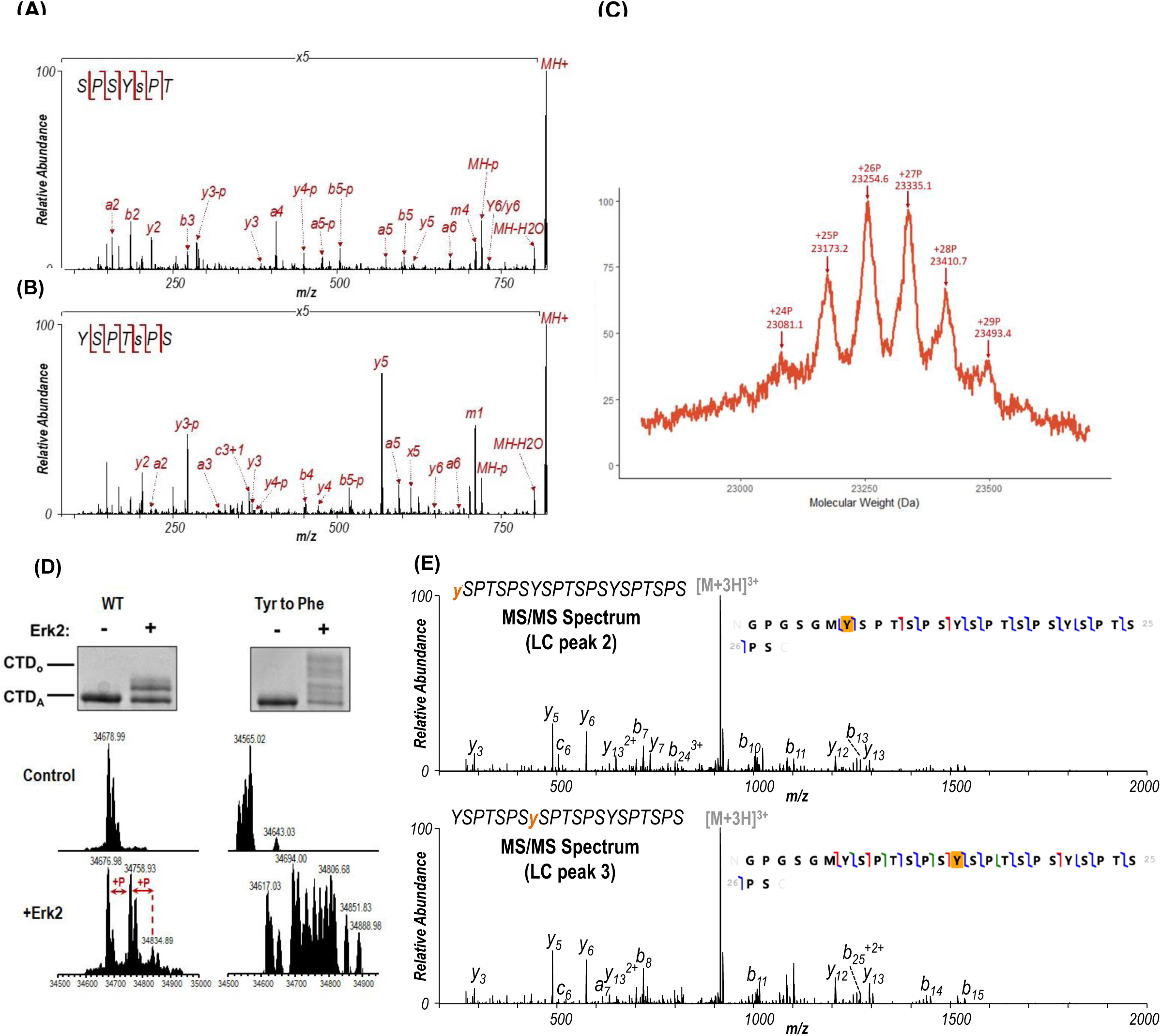
Mass spectrometry analysis of yCTD treated by kinases. Related to Figure 1 **and** Figure 2. (A, B) LC-UVPD-MS/MS MS2 spectra of yCTD treated with P-TEFb: (A) SPSYsPT and (B) YSPTsPS. These spectra correspond to the LC traces in Figure 1A. The locations of the confirmed phosphorylation sites are indicated as lowercase y or s in the sequences. (C) MALDI-TOF of yCTD treated with P-TEFb, zoomed view to show detail. Portion of MALDI-MS spectra of 3C-protease digested yCTD constructs treated with P-TEFb. Mass labels indicate m/z at the various peak maxima. Arrows indicate the maximum and minimum m/z peaks for kinase treatment sample. “+#P” notation indicates an approximate number of phosphates added based on mass shifts relative to no kinase control. (D) EMSA to show the effect of YtoF mutation on CTD gel shift upon kinase phosphorylation. EMSA and intact mass analysis result for Drosophila CTD 9mer before and after treatment with Erk2. (E) LC-UVPD-MS/MS spectra of 3CTD treated with c-Abl. Diagnostic fragmentation of mon-phosphorylated peptides corresponding to LC peaks 2 and 3 from Figure 2B are shown in the top and bottom spectra, respectively

**Supplementary Figure S3.**
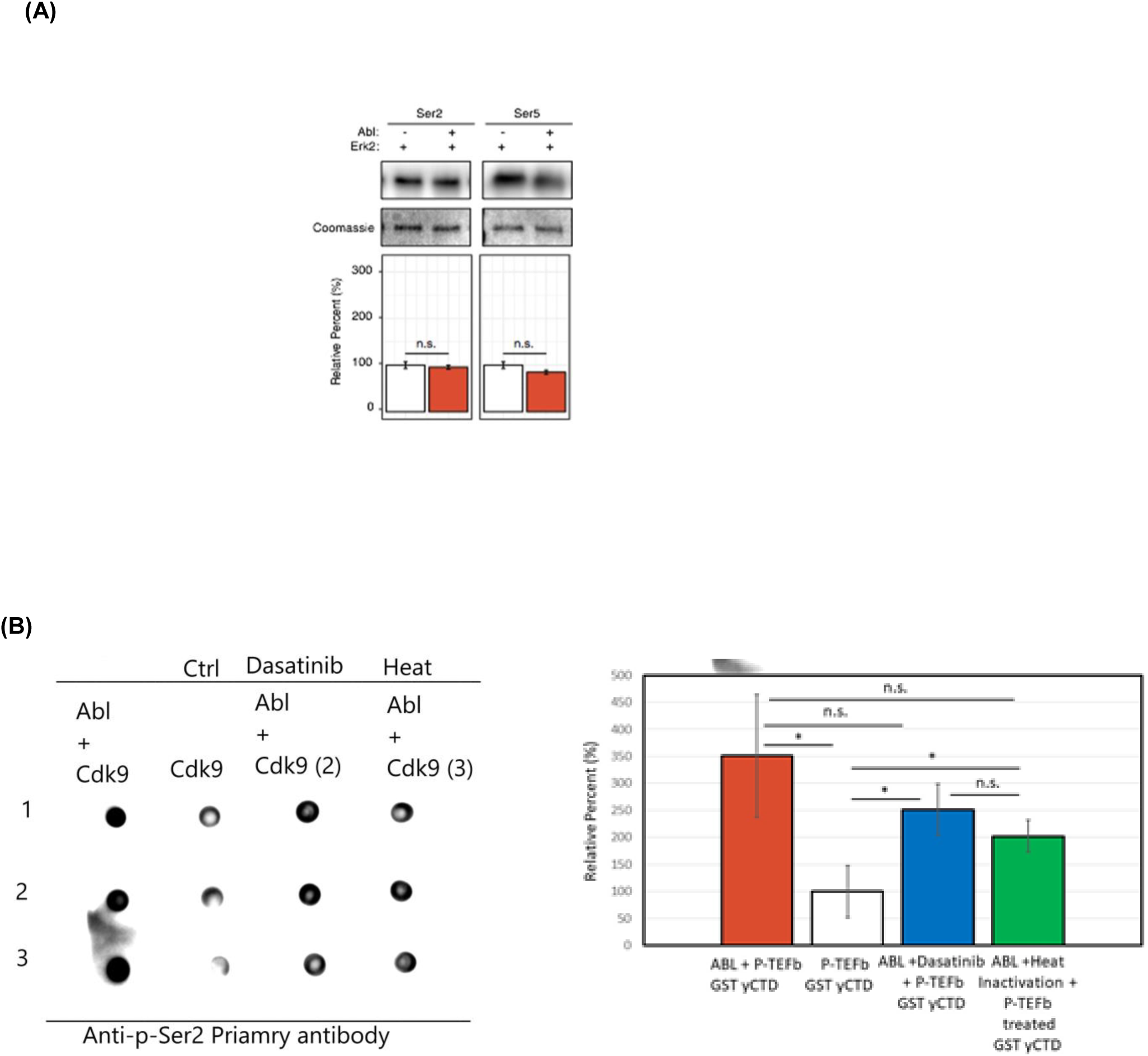
Phosphorylation of different CTD constructs by CTD kinases, related to Figure 3. (A) Western blot analysis of yCTD treated in tandem with c-Abl and Erk2. Representative images (top) and quantification (bottom) of western blot analysis of yeast CTD treated with Erk2 alone (left, white) and tandemly with c-Abl followed by Erk2 (right, red). Tandem treatment of c-Abl followed by Erk2 does not significantly alter the epitope abundance of phosphorylated Ser2 or Ser5. Significance determined by Welch’s t-test (n.s. = not significant (p-value > 0.05)), n = 3, error bars indicate SEM. (B) Dot blot analysis of the tandem treatment of c-Abl and P-TEFb. c-Abl treated GST yCTD followed by Dasatinib inhibition before P-TEFb treatment (blue), c-Abl treated GST yCTD followed by heat inactivation of c-Abl before treatment with P-TEFb (green) with corresponding P-TEFb only treated GST yCTD (white) and c-Abl followed by P-TEFb treated GST yCTD (red). Significance determined by the t-test (* = p-value < 0.05) n = 3, n.s. = not significant (p-value > 0.05), n = 3, error bars indicate SEM from three technical replicates.

**Supplementary Figure S4.**
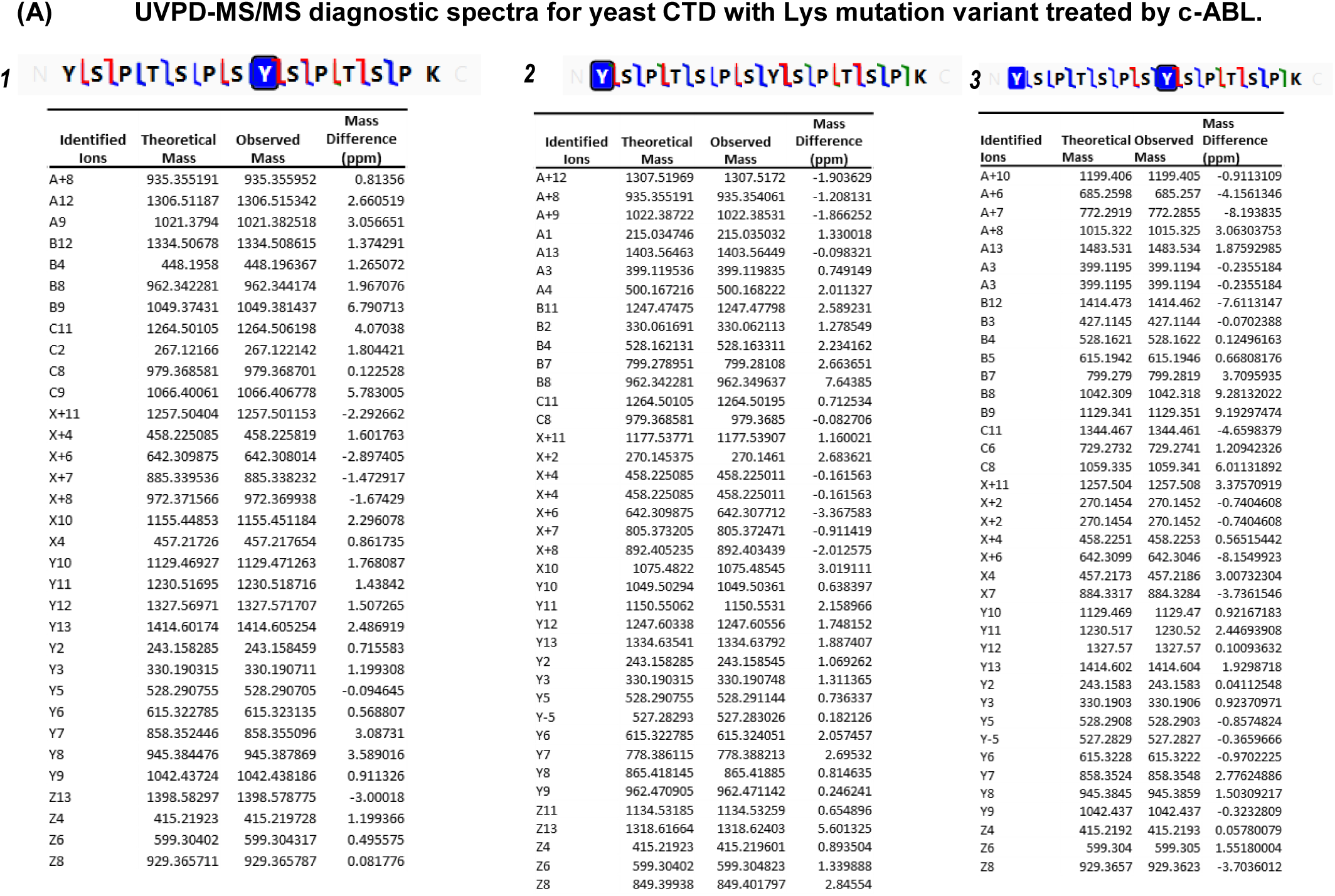
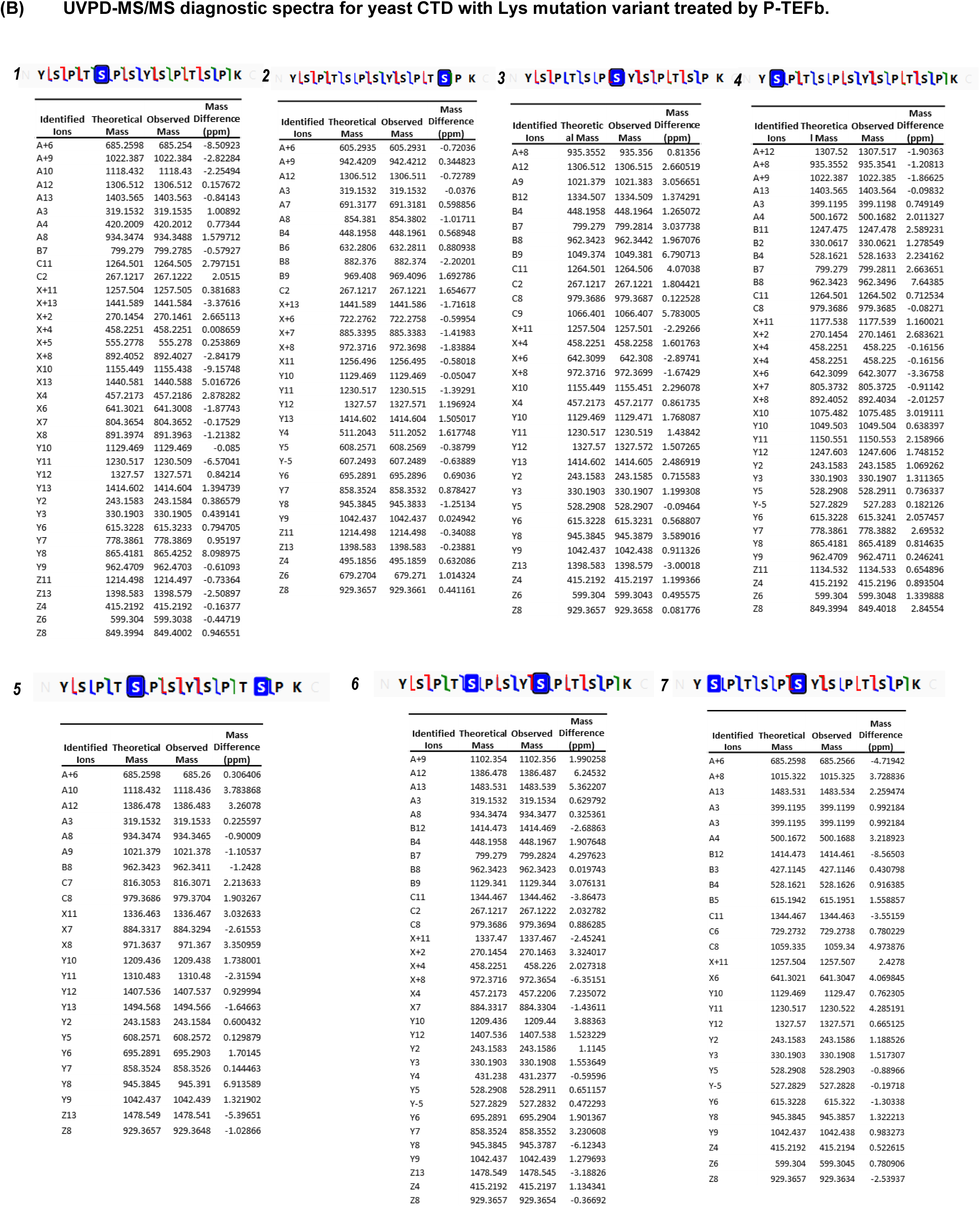
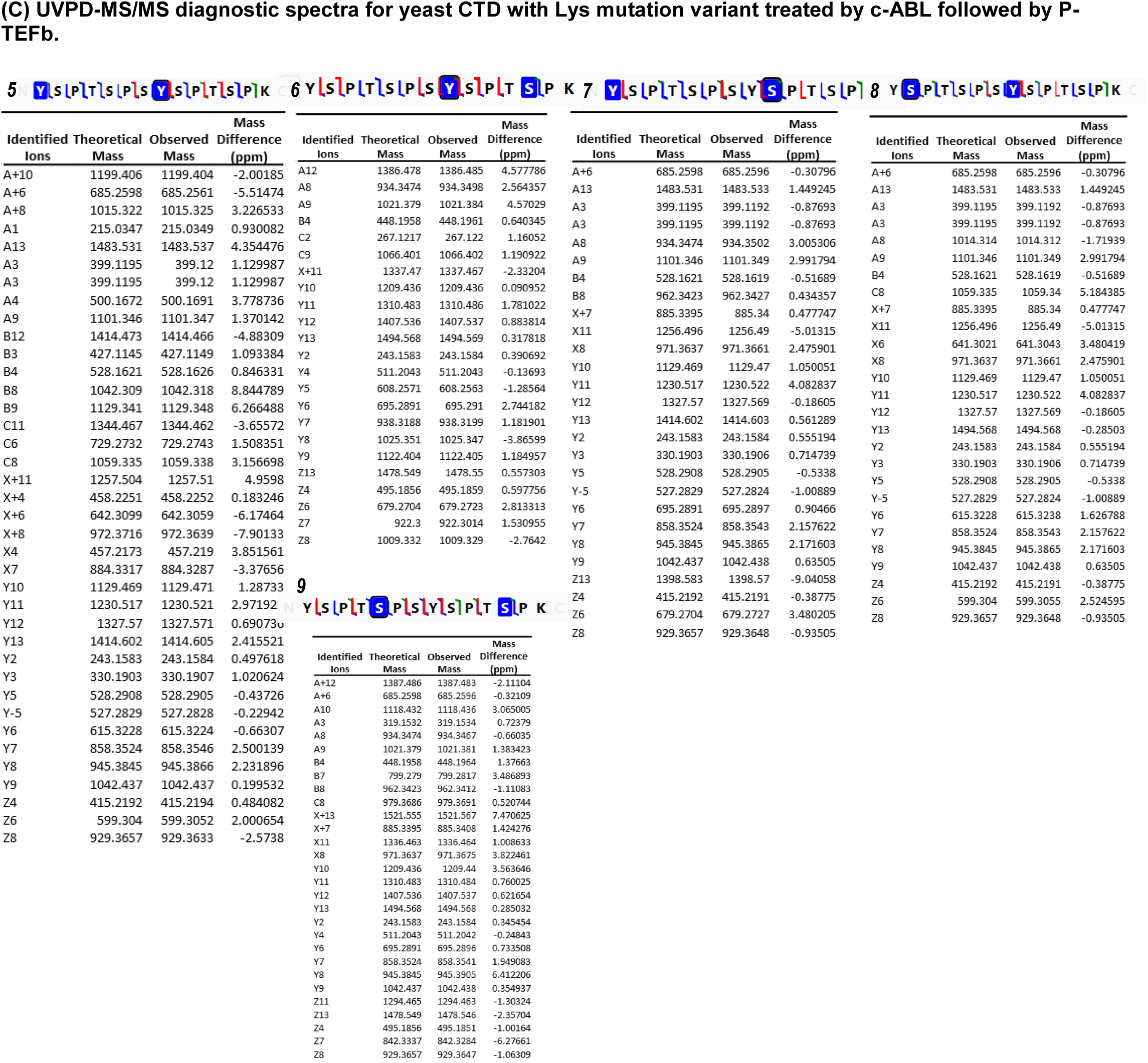
LC-UVPD-mass spectra yCTD-Lys treated with CTD kinases. Related to Figure 3. The tables show the fragments matched from UVPD spectra. The confirmed phosphorylated residues are highlighted with blue boxes in the sequence, and backbone cleavages that produce diagnostic fragment ions are designated by color-coded slash marks (a/x green, b/y blue, c/z red) in the sequence.

**Supplementary Figure S5.**
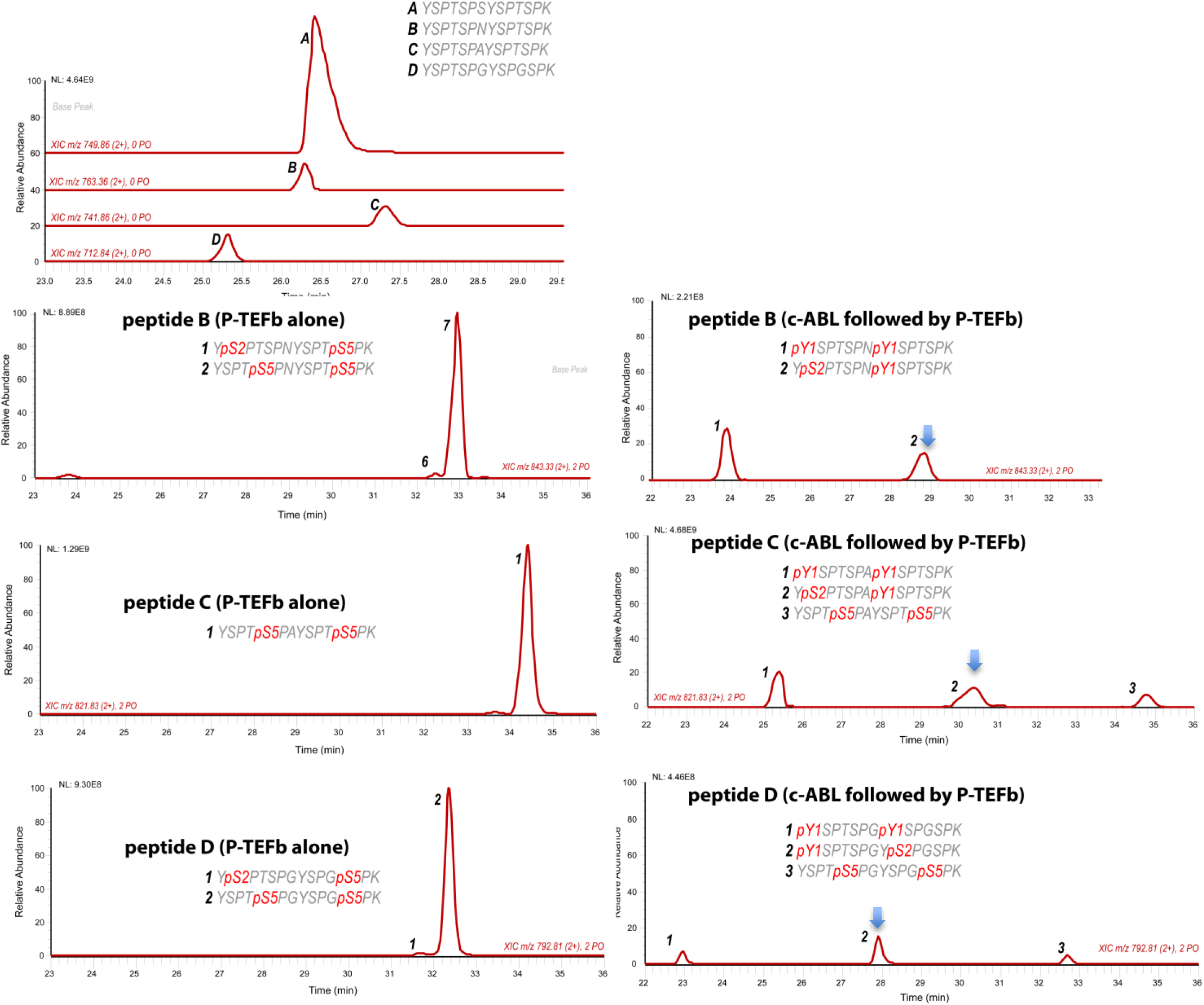
LC-UVPD-mass spectra yCTD-Lys treated with CTD kinases. Related to Figure 3. LC-UVPD-MS/MS analysis of yeast CTD with inserted Lys in every other repeat treated with c-Abl showing extracted ion chromatograms. For claritiy, only doubly phosphorylated species (m/z 792.81, 2+ charge state) peptides are shown. Peptide A is most abundant and its spectra are shown in Figure 3G-H. The less abundant peptides (peptide B, C, D) are shown here when different kinase treatment was applied. For the clarity, only doubly phosphorylation species were shown. The phoshoryl-species containing both pTyr1 and pSer2 are highlighted with arrows.

**Supplementary Figure 6.**
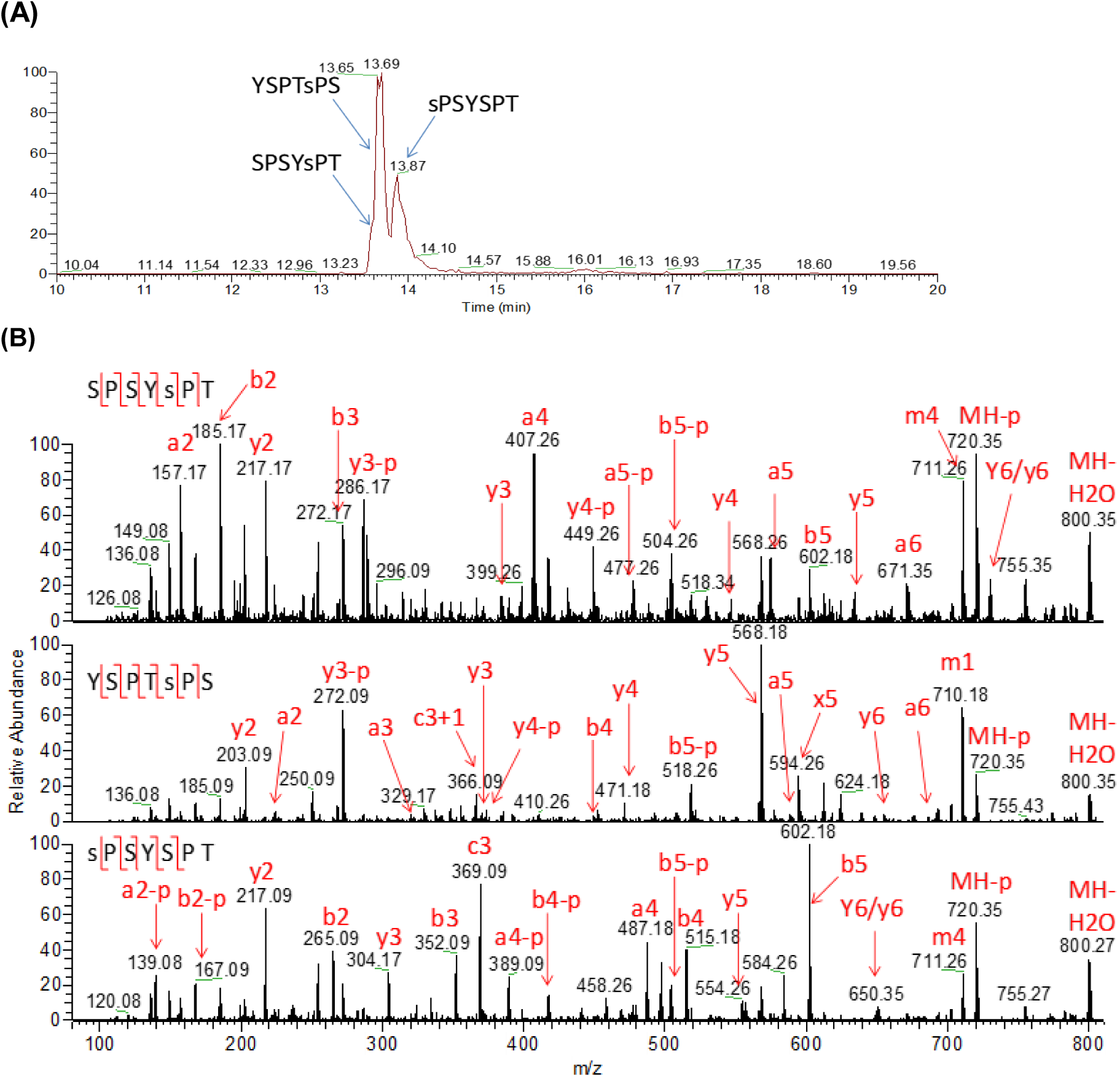
Analysis of yCTD treated in tandem with TFIIH and P-TEFb. Related to Figure 3. (A) Extracted ion chromatogram of mono-phosphorylated heptad (m/z 818, 2+ charge state) peptides. (B) UVPD mass spectra of three mono-phosphorylated heptads. The locations of the confirmed phosphorylation sites are indicated as lowercase y or s in the sequences.

**Figure S7:**
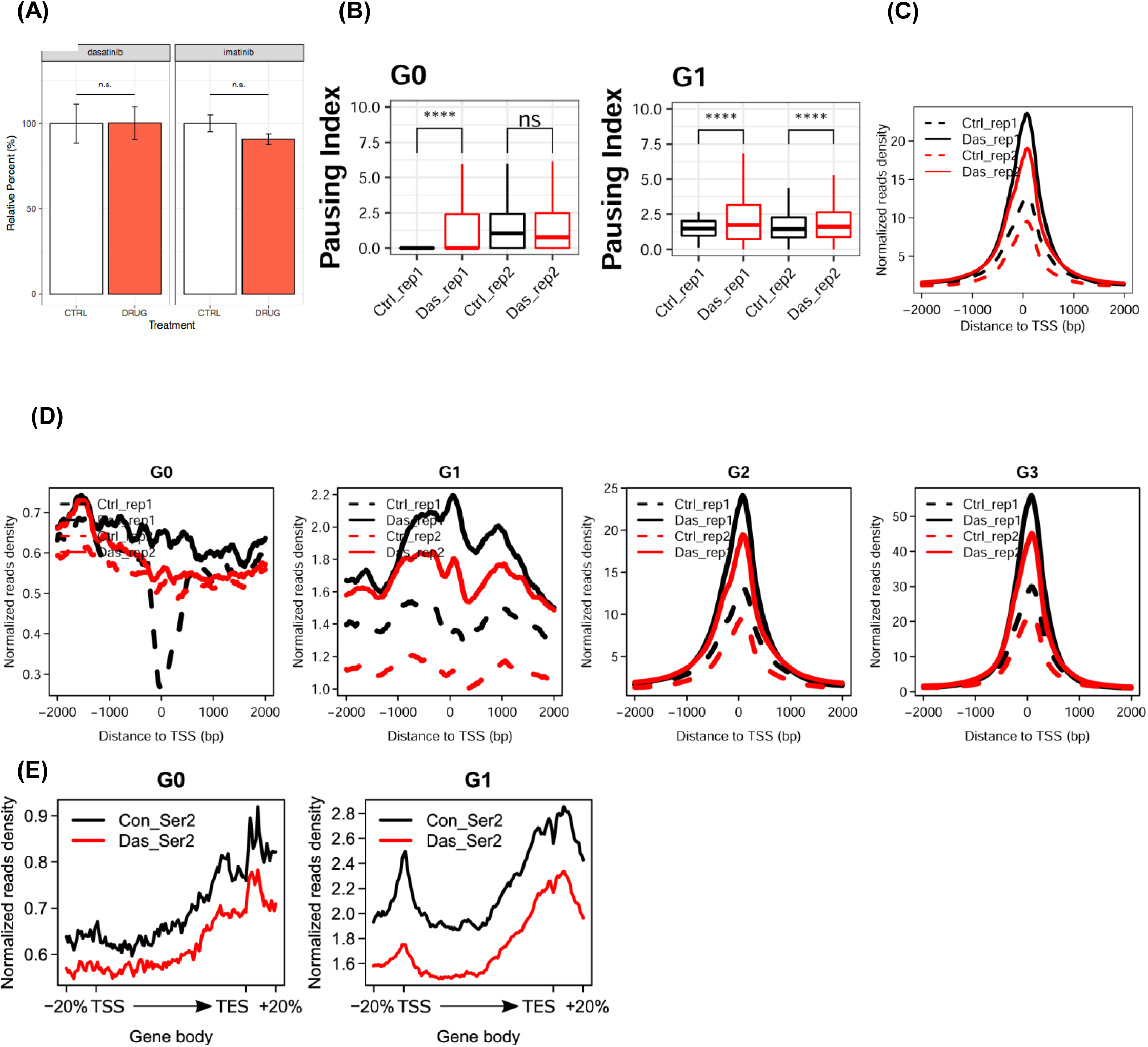
Cell-based analysis of phosphorylation of RNA polymerase II. Related to Figure 4-5. (A) Quantification of POLR2C western blot from Figure 4. POLR2C bands from dasatinib and imatinib blots (indicated) from Figure 4 were quantified. Dasatinib treatment POLR2C quantification was performed on the same blots providing the Ser2 and Ser5 data. Imatinib treatment POLR2C quantification was performed on the same blots providing POLR2A and Ser5 data. No significant change in POLR2C levels was observed for any blot analyzed. Epitope signals normalized against tubulin loading control. Significance determined by Welch’s t-test (n.s. = not significant (p-value > 0.05)), error bars indicate SEM, n=6). (B) Boxplot of pausing index changes of HEK293 cells on the genes from G0 and G1 clusters upon Tyr1 phosphorylation. “ns” indicates a p-value higher than 0.05. “****” indicates p-value less than 0.0001. (C) Lineplot of pausing indexes on all genes. Solid and dotted lines are representing two biological replicate. (D) Lineplot of pausing indexes on genes for each cluster. (E) The profiling of pSer2 signal along gene body under control and dasatinib treatment for genes from cluster G0 and G1.

## References

Bartkowiak, B., Liu, P., Phatnani, H.P., Fuda, N.J., Cooper, J.J., Price, D.H., Adelman, K., Lis, J.T., and Greenleaf, A.L. (2010). CDK12 is a transcription elongation-associated CTD kinase, the metazoan ortholog of yeast Ctk1. Genes Dev 24, 2303–2316.

Baskaran, R., Chiang, G.G., Mysliwiec, T., Kruh, G.D., and Wang, J.Y. (1997). Tyrosine phosphorylation of RNA polymerase II carboxyl-terminal domain by the Abl-related gene product. J Biol Chem 272, 18905–18909.

Baskaran, R., Dahmus, M.E., and Wang, J.Y. (1993). Tyrosine phosphorylation of mammalian RNA polymerase II carboxyl-terminal domain. Proc Natl Acad Sci U S A 90, 11167–11171.

Baskaran, R., Escobar, S.R., and Wang, J.Y. (1999). Nuclear c-Abl is a COOH-terminal repeated domain (CTD)-tyrosine (CTD)-tyrosine kinase-specific for the mammalian RNA polymerase II: possible role in transcription elongation. Cell Growth Differ 10, 387–396.

Bataille, A.R., Jeronimo, C., Jacques, P.E., Laramee, L., Fortin, M.E., Forest, A., Bergeron, M., Hanes, S.D., and Robert, F. (2012). A universal RNA polymerase II CTD cycle is orchestrated by complex interplays between kinase, phosphatase, and isomerase enzymes along genes. Mol Cell 45, 158–170.

Buratowski, S. (2003). The CTD code. Nat Struct Biol 10, 679–680.

Burger, K., Schlackow, M., and Gullerova, M. (2019). Tyrosine kinase c-Abl couples RNA polymerase II transcription to DNA double-strand breaks. Nucleic Acids Res.

Chao, S.H., and Price, D.H. (2001). Flavopiridol inactivates P-TEFb and blocks most RNA polymerase II transcription in vivo. J Biol Chem 276, 31793–31799.

Chapman, R.D., Heidemann, M., Hintermair, C., and Eick, D. (2008). Molecular evolution of the RNA polymerase II CTD. Trends Genet 24, 289–296.

Chen, H.H., Wong, Y.H., Geneviere, A.M., and Fann, M.J. (2007). CDK13/CDC2L5 interacts with L-type cyclins and regulates alternative splicing. Biochem Biophys Res Commun 354, 735–740.

Corden, J.L. (2013). RNA polymerase II C-terminal domain: Tethering transcription to transcript and template. Chem Rev 113, 8423–8455.

Czudnochowski, N., Bosken, C.A., and Geyer, M. (2012). Serine-7 but not serine-5 phosphorylation primes RNA polymerase II CTD for P-TEFb recognition. Nat Commun 3, 842.

Descostes, N., Heidemann, M., Spinelli, L., Schuller, R., Maqbool, M.A., Fenouil, R., Koch, F., Innocenti, C., Gut, M., Gut, I., et al. (2014). Tyrosine phosphorylation of RNA polymerase II CTD is associated with antisense promoter transcription and active enhancers in mammalian cells. Elife 3, e02105.

Ebmeier, C.C., Erickson, B., Allen, B.L., Allen, M.A., Kim, H., Fong, N., Jacobsen, J.R., Liang, K., Shilatifard, A., Dowell, R.D., et al. (2017). Human TFIIH Kinase CDK7 Regulates Transcription-Associated Chromatin Modifications. Cell Rep 20, 1173–1186.

Eick, D., and Geyer, M. (2013). The RNA polymerase II carboxy-terminal domain (CTD) code. Chem Rev 113, 8456–8490.

Feaver, W.J., Svejstrup, J.Q., Henry, N.L., and Kornberg, R.D. (1994). Relationship of CDK-activating kinase and RNA polymerase II CTD kinase TFIIH/TFIIK. Cell 79, 1103–1109.

Fellers, R.T., Greer, J.B., Early, B.P., Yu, X., LeDuc, R.D., Kelleher, N.L., and Thomas, P.M. (2015). ProSight Lite: graphical software to analyze top-down mass spectrometry data. Proteomics 15, 1235–1238.

Franco, L.C., Morales, F., Boffo, S., and Giordano, A. (2018). CDK9: A key player in cancer and other diseases. J Cell Biochem 119, 1273–1284.

Gardner, M.W., Vasicek, L.A., Shabbir, S., Anslyn, E.V., and Brodbelt, J.S. (2008). Chromogenic cross-linker for the characterization of protein structure by infrared multiphoton dissociation mass spectrometry. Anal Chem 80, 4807–4819.

Hamilton, N. (2015). smoother: Functions Relating to the Smoothing of Numerical Data.

Harlen, K.M., and Churchman, L.S. (2017). The code and beyond: transcription regulation by the RNA polymerase II carboxy-terminal domain. Nat Rev Mol Cell Biol 18, 263–273.

Heidemann, M., Hintermair, C., Voss, K., and Eick, D. (2013). Dynamic phosphorylation patterns of RNA polymerase II CTD during transcription. Biochim Biophys Acta 1829, 55–62.

Hsin, J.P., Li, W., Hoque, M., Tian, B., and Manley, J.L. (2014). RNAP II CTD tyrosine 1 performs diverse functions in vertebrate cells. Elife 3, e02112.

Jeronimo, C., Bataille, A.R., and Robert, F. (2013). The writers, readers, and functions of the RNA polymerase II C-terminal domain code. Chem Rev 113, 8491–8522.

Jeronimo, C., Collin, P., and Robert, F. (2016). The RNA Polymerase II CTD: The Increasing Complexity of a Low-Complexity Protein Domain. J Mol Biol 428, 2607–2622.

Khokhlatchev, A., Xu, S., English, J., Wu, P., Schaefer, E., and Cobb, M.H. (1997). Reconstitution of mitogen-activated protein kinase phosphorylation cascades in bacteria. Efficient synthesis of active protein kinases. J Biol Chem 272, 11057–11062.

Klein, D.R., Holden, D.D., and Brodbelt, J.S. (2016). Shotgun Analysis of Rough-Type Lipopolysaccharides Using Ultraviolet Photodissociation Mass Spectrometry. Anal Chem 88, 1044–1051.

Knight, G.W., and McLellan, D. (2004). Use and limitations of imatinib mesylate (Glivec), a selective inhibitor of the tyrosine kinase Abl transcript in the treatment of chronic myeloid leukaemia. Br J Biomed Sci 61, 103–111.

Langmead, B., Trapnell, C., Pop, M., and Salzberg, S.L. (2009). Ultrafast and memory-efficient alignment of short DNA sequences to the human genome. Genome Biol 10, R25.

Madsen, J.A., Boutz, D.R., and Brodbelt, J.S. (2010). Ultrafast ultraviolet photodissociation at 193 nm and its applicability to proteomic workflows. Journal of proteome research 9, 4205–4214.

Marshall, N.F., Peng, J., Xie, Z., and Price, D.H. (1996). Control of RNA polymerase II elongation potential by a novel carboxyl-terminal domain kinase. J Biol Chem 271, 27176–27183.

Mayer, A., Heidemann, M., Lidschreiber, M., Schreieck, A., Sun, M., Hintermair, C., Kremmer, E., Eick, D., and Cramer, P. (2012). CTD tyrosine phosphorylation impairs termination factor recruitment to RNA polymerase II. Science 336, 1723–1725.

Mayfield, J.E., Robinson, M.R., Cotham, V.C., Irani, S., Matthews, W.L., Ram, A., Gilmour, D.S., Cannon, J.R., Zhang, Y.J., and Brodbelt, J.S. (2017). Mapping the Phosphorylation Pattern of Drosophila melanogaster RNA Polymerase II Carboxyl-Terminal Domain Using Ultraviolet Photodissociation Mass Spectrometry. ACS Chem Biol 12, 153–162.

Ni, Z., Schwartz, B.E., Werner, J., Suarez, J.R., and Lis, J.T. (2004). Coordination of transcription, RNA processing, and surveillance by P-TEFb kinase on heat shock genes. Mol Cell 13, 55–65.

Portz, B., Lu, F., Gibbs, E.B., Mayfield, J.E., Rachel Mehaffey, M., Zhang, Y.J., Brodbelt, J.S., Showalter, S.A., and Gilmour, D.S. (2017). Structural heterogeneity in the intrinsically disordered RNA polymerase II C-terminal domain. Nat Commun 8, 15231.

Rahl, P.B., Lin, C.Y., Seila, A.C., Flynn, R.A., McCuine, S., Burge, C.B., Sharp, P.A., and Young, R.A. (2010). c-Myc regulates transcriptional pause release. Cell 141, 432–445.

Robinson, J.T., Thorvaldsdottir, H., Winckler, W., Guttman, M., Lander, E.S., Getz, G., and Mesirov, J.P. (2011). Integrative genomics viewer. Nat Biotechnol 29, 24–26.

Schneider, C.A., Rasband, W.S., and Eliceiri, K.W. (2012). NIH Image to ImageJ: 25 years of image analysis. Nat Methods 9, 671–675.

Schuller, R., Forne, I., Straub, T., Schreieck, A., Texier, Y., Shah, N., Decker, T.M., Cramer, P., Imhof, A., and Eick, D. (2016). Heptad-Specific Phosphorylation of RNA Polymerase II CTD. Mol Cell 61, 305–314.

Suh, H., Ficarro, S.B., Kang, U.B., Chun, Y., Marto, J.A., and Buratowski, S. (2016). Direct Analysis of Phosphorylation Sites on the Rpb1 C-Terminal Domain of RNA Polymerase II. Mol Cell 61, 297–304.

Team, R.C. (2017). R: A language and environment for statistical computing, R.F.f.S. Computing, ed. (Vienna, Austria).

Tee, W.W., Shen, S.S., Oksuz, O., Narendra, V., and Reinberg, D. (2014). Erk1/2 activity promotes chromatin features and RNAPII phosphorylation at developmental promoters in mouse ESCs. Cell 156, 678–690.

Wada, T., Takagi, T., Yamaguchi, Y., Watanabe, D., and Handa, H. (1998). Evidence that P-TEFb alleviates the negative effect of DSIF on RNA polymerase II-dependent transcription in vitro. EMBO J 17, 7395–7403.

West, M.L., and Corden, J.L. (1995). Construction and analysis of yeast RNA polymerase II CTD deletion and substitution mutations. Genetics 140, 1223–1233.

Wickhan, H. (2009). ggplot2: Elegant Graphics for Data Analysis (New York: Springer-Verlag).

Winter, G.E., Rix, U., Carlson, S.M., Gleixner, K.V., Grebien, F., Gridling, M., Muller, A.C., Breitwieser, F.P., Bilban, M., Colinge, J., et al. (2012). Systems-pharmacology dissection of a drug synergy in imatinib-resistant CML. Nat Chem Biol 8, 905–912.

Zeitlinger, J., Stark, A., Kellis, M., Hong, J.W., Nechaev, S., Adelman, K., Levine, M., and Young, R.A. (2007). RNA polymerase stalling at developmental control genes in the Drosophila melanogaster embryo. Nat Genet 39, 1512–1516.

Zhang, Y., Liu, T., Meyer, C.A., Eeckhoute, J., Johnson, D.S., Bernstein, B.E., Nusbaum, C., Myers, R.M., Brown, M., Li, W., et al. (2008). Model-based analysis of ChIP-Seq (MACS). Genome Biol 9, R137.

